# Species and strain diversity in *Staphylococcus* drive divergent host responses in human skin

**DOI:** 10.64898/2026.04.30.720712

**Authors:** Ruoyu Yang, Morgan M. Severn, Elizabeth S. Aiken, Wei Zhou, Anita Y. Voigt, Gabrielle M. Walker, Aaron Y. Koh, Minghao Gong, Maheshwor Thapa, Shuzhao Li, Leonard M. Milstone, Julia Oh

## Abstract

The skin microbiome regulates key skin processes, yet the functional diversity of a dominant genus, *Staphylococcus,* remains poorly resolved at the strain level for multiple species across its pathogenic and commensal continuum. It is likely that *Staphylococcus’* effects on skin are diverse at these finest taxonomic resolutions, but current skin models lack the physiological relevance and scalability needed to profile this diversity. Using an organotypic 3D human skin model (reconstructed human epidermis, RHE), we profiled skin responses to 187 *Staphylococcus* strains across seven dominant species. Canonically ‘pathogenic’ species (e.g., *S. aureus*) induced broad inflammatory responses, whereas prototypical ‘commensal’ species (e.g., *S. hominis*) elicited more nuanced effects on innate immune and skin barrier responses. Strikingly, *S. epidermidis* displayed pronounced strain-level heterogeneity, with subsets inducing either ‘commensal’ or ‘pathogen’-like responses despite lacking canonical virulence factors, suggesting pleiotropic effects. Comparative genomics, dual-transcriptomics, untargeted metabolomics, and growth phenotyping revealed species- and strain-specific traits underlying these differential effects on RHE, including the presence of select cell surface proteins and differential arginine metabolism. Together, our study provides the first high-throughput, species- and strain-resolved analysis of skin–*Staphylococcus* interactions, offering mechanistic insights and a platform for microbiome-based strategies to modulate skin inflammation and diseases.

**One-line summary:** High-throughput profiling of *Staphylococcus* in a human skin model shows that species- and strain-level diversity underlies a continuum of host barrier and immune responses.

## INTRODUCTION

The skin microbiome is a key regulator of skin physiology, with *Staphylococcus* (*S.*) species serving as keystone and influential colonizers. Commensal staphylococci such as *S. epidermidis* contribute to skin processes by modulating skin immunity^1–3^, barrier function^4–6^, and skin-microbiome homeostasis^7–13^. In contrast, prototypically pathogenic species such as *S. aureus* drive inflammation and contribute to numerous skin diseases^14^. However, this pathogen-commensal dichotomy has blurred as *Staphylococcus-*host interactions are increasingly identified as strain-specific, with interspecies and intraspecies variation attributed to both beneficial and disease-promoting traits. For example, some *S. epidermidis* strains enhance skin innate immunity and protection against cutaneous pathogens in mice^3^; while, others have been linked to disease flares in atopic dermatitis^15^.

We previously defined the extraordinary strain-level genetic diversity of *S. epidermidis*, where hundreds of genes could differ between strains isolated from the same skin site of a single individual. This diversity conferred unique phenotypes, including antibiotic resistance and the ability to produce potent antimicrobials^16,17^. These studies very likely portend a similar diversity in other *Staphylococcus* species, whose prevalence have recently come to light via metagenomic surveys of the skin microbiome^18–23^.

However, the contributions of such inter- and intra-species genetic diversity, of both *S. epidermidis* and other *Staphylococcus* species, to skin processes remain poorly understood. A major barrier to addressing this question is the lack of scalable and physiologically relevant systems; the natural microbial diversity of human skin cannot be controlled for, while the ability to control diversity in germ-free mice is countered by the physiological differences between mouse and human skin. Moreover, neither *ex vivo* human skin explants nor germ-free mice can be efficiently scaled for studies of hundreds of microbial isolates, and screens in 2D submerged skin cell cultures can have limited physiological relevance.

Here, we used a 3D human skin model, reconstructed human epidermis (RHE), to examine responses of stratified keratinocytes to colonization by *Staphylococcus*. Keratinocytes are the predominant cell type in the epidermis and thereby central mediators in sensing and responding to skin-colonizing microorganisms via an array of receptors such as Toll-like receptors (TLRs)^24^ and the aryl hydrocarbon receptor (AhR)^25^. Keratinocytes actively regulate colonization of *Staphylococcus* by modulating bacterial adhesion to the outermost layer of the epidermis, the stratum corneum, and by secreting antimicrobial peptides and proinflammatory cytokines, which coordinate the recruitment of immune cells to sites of colonization^26^ or infection of *S. aureus*^27^. Notably, RHE’s stratified keratinocytes at air liquid interface better mimic the skin’s epidermal structure in contrast to submerged keratinocyte cultures. Here, its use allowed us to achieve high-throughput discovery of primary interactions between microbes and keratinocytes while preserving the inherently structured nature of this host-microbiome interaction.

We hypothesized that genetically diverse strains across *Staphylococcus* species differentially affect skin processes in the human epidermis. We colonized RHE with 187 strains from seven dominant species of *Staphylococcus* and examined the transcriptional responses of the RHE after 18 hours. We performed comparative genomics, untargeted metabolomics, and defined phenotypes relevant to skin colonization *in vitro* to decipher the traits that underlie these differential effects on RHE.

Our study reveals remarkable species- and strain-level heterogeneity in how *Staphylococcus* shapes human epidermal responses. While *S. aureus, S. haemolyticus,* and *S. lugdunensis* drove robust proinflammatory ‘pathogenic’ responses, *S. hominis, S. capitis,* and *S. pasteuri* elicited comparatively attenuated ‘commensal’-like responses. Notably, *S. epidermidis* strains separated into two predominant phenotypes in which a part elicited a more ‘pathogenic’ effect with respect to inflammatory and skin barrier processes, with the remaining strains more closely resembling the other ‘commensal’ species. An integrated multi-omics analysis of genomes, staphylococcal transcriptomics, and functional analyses identified candidate drivers of these phenotypes, including variation in cell surface factors, arginine metabolism, and biofilm and blood growth traits enriched in these more ‘pathogenic’ *S. epidermidis*.

Taken together, these findings establish a scalable, physiologically relevant framework for dissecting host-microbiome interactions at strain resolution and provide a foundation for understanding the mechanisms by which these keystone members of the skin microbiome modulate skin barrier and immune responses in health and disease.

## RESULTS

### Data characteristics and analysis overview

We selected 187 strains from seven species from our biorepository of ∼5000 sequenced strains of *Staphylococcus* isolated from healthy human skin, congenital ichthyosis, and miscellaneous other human sources^28–30^ (Table S1). Strains were selected to maximize genetic diversity of stereotypical ‘commensal’ (n=50 each from *S. epidermidis*, *S. capitis*, *S. hominis,* n=7 for *S. pasteuri)* and ‘pathogenic’ species (n=10 each from *S. aureus*, *S. haemolyticus*, *S. lugdunensis*); hereafter for simplicity, we will use these designations though recognizing the reality of a continuous phenotypic spectrum. Each strain was colonized apically onto RHE (n=3 per strain) for 18 hours, then harvested for dual RNA-seq to associate the host and staphylococcal transcriptome. In parallel, we performed untargeted metabolomics on the supernatant and lysate of each strain to capture the metabolic diversity. Comparative genomics further identified potential genes underlying phenotypic differences, and finally, we generated select experimental data to validate our findings. The overall experimental and analytic schematics are shown in Figures 1A and S1A.

**Figure 1.**
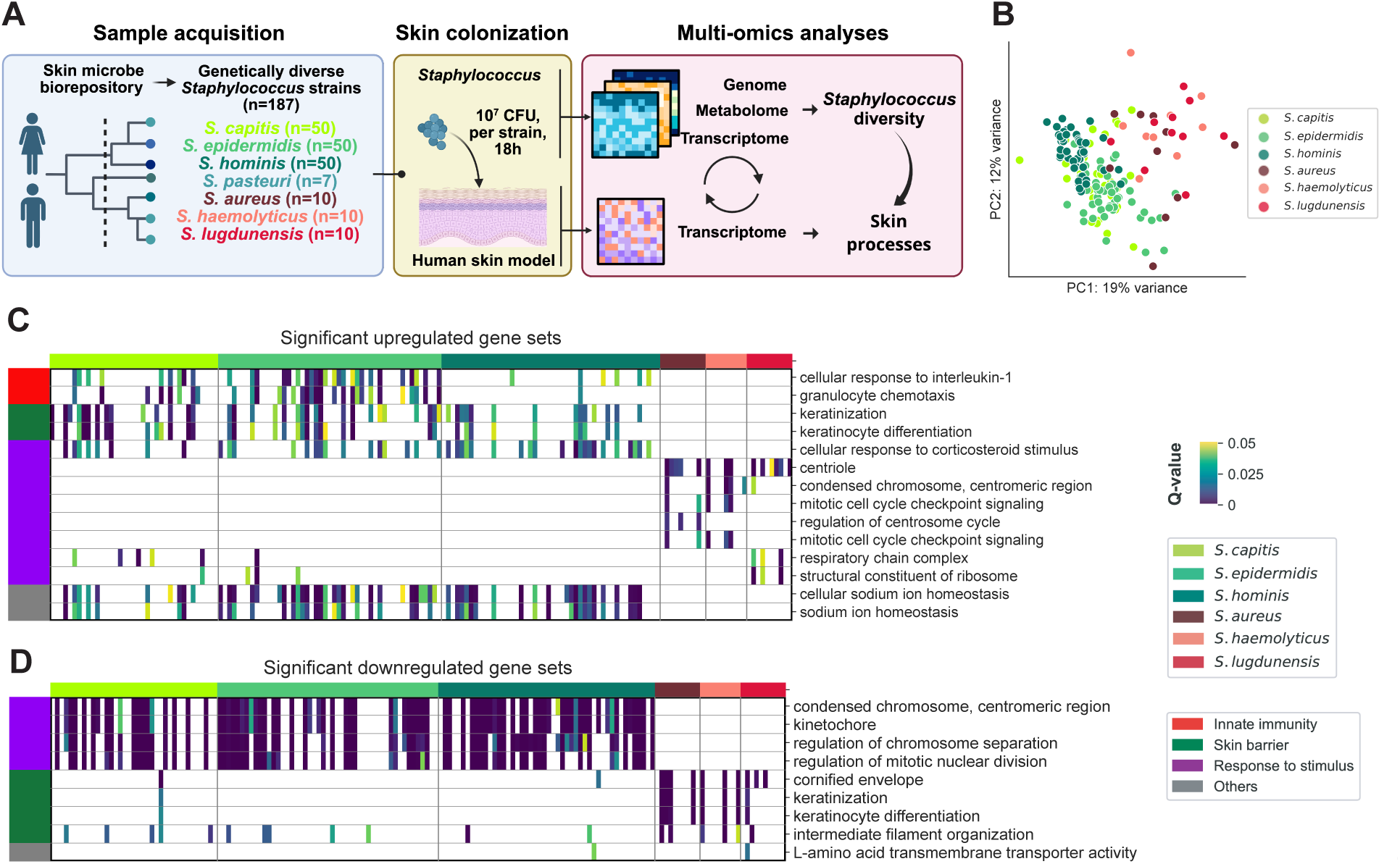
*Staphylococcus* colonization drives diverse transcriptional programs in reconstructed human epidermis (RHE) **(A)** Scheme of study design. 187 strains from 7 *Staphylococcus* (*S.*) species were selected from our biorepository by maximal genetic diversity as defined by core genome phylogeny. Each strain was topically colonized at 10^7^ CFU in water onto a human 3D skin model comprised of differentiated keratinocytes (reconstructed human epidermis, RHE) for 18 hours, after which total RNA was extracted for dual RNA-seq. We performed complementary ‘omics analyses to determine the *Staphylococcus* and RHE transcriptional responses to colonization and to identify potential microbial mediators underlying differential effects of staphylococcal strains on RHE skin processes. **(B)** Principal components analysis (PCA) of genome-wide transcriptional responses of RHE following *Staphylococcus* colonization (n=3 replicates per strain; n=14 controls), based on log_2_ fold-change (log2FC). Points are colored by species. **(C-D)** Heatmaps of enriched Gene Ontology (GO) gene sets among upregulated (C) and downregulated genes in *Staphylococcus-*colonized RHE vs. vehicle controls. Gene Set Enrichment Analysis (GSEA) on differentially expressed genes (DEGs, if FDR adjusted P-value ≤ 0.05 and |log2FC| ≥ 1) is shown for each strain. Rows represent GO gene sets and columns *Staphylococcus* strains. The union of the top three gene sets per species (ranked by mean gene ratio) is shown. Cells are colored by Q-value (FDR adjusted P-value) if significant (Q ≤ 0.05); non-significant entries are blank.

### RHE transcriptional profiling reveals diverse responses to *Staphylococcus* species and *S. epidermidis* strains

#### Global transcriptional effects

We first examined the global RHE transcriptional response to colonization. Principal component (PC) analysis showed distinct species-level differences between stereotypical commensal vs. pathogenic species, primarily explained by PC1. Interestingly, strain variation (captured by PC2) were also evident for all species, especially *S. epidermidis* (Figure 1B). Differentially expressed genes (DEGs) in RHE colonized by stereotypical commensals vs. vehicle control (n=14) were markedly fewer than for pathogenic species. Only ∼10% of DEGs were shared across all species, supporting distinct species-level effects of *Staphylococcus* colonization. Within species, ∼30-60% of DEGs overlapped across strains while others were unique, indicating substantial intraspecies heterogeneity (Table S2). Functionally, commensal strains significantly upregulated Gene Ontology (GO) gene sets associated with innate immunity (e.g., cellular response to interleukin-1) and skin barrier function (e.g., keratinization), while pathogenic strains upregulated gene sets related to response to stimulus and downregulated pathways linked to skin barrier functions (e.g., cornified envelope) (Figures 1C-1D). Given these distinct effects on innate immunity and barrier-associated pathways, we focused subsequent analyses on how *Staphylococcus* modulate these key skin processes in RHE.

#### Innate immune effects

To more closely examine *Staphylococcus* effects on skin immunity, we curated a panel of 142 related genes (Table S2). To further reduce dimensionality, genes were grouped into modules based on hierarchical clusters, akin to GO gene sets. As anticipated, stereotypical pathogens elicited strong proinflammatory responses in RHE characterized by strong upregulation of innate immunity module (M)2 genes (Figure 2A). Strikingly, the effects of strains of commensal species segregated into two major clusters, which we termed “commensals” vs. “inflammatory commensals”. Commensals, comprised mainly of *S. capitis* and *S. hominis* and ∼1/3 of the S*. epidermidis* strains, induced only modest innate immune responses. In contrast, inflammatory commensals, comprised mainly of *S. epidermidis* strains, elicited transcriptional profiles that more closely resembled the pathogen-colonized RHE, with strong upregulation of innate immunity M2 and M3 genes, which included proinflammatory chemokines (*CXCL8* and *CCL20*), cytokines (e.g., *IL1A*, *IL23A* and *IL36G*), regulatory factors (e.g., *NFKB2*), and antimicrobial peptides (e.g., *DEFB4A* and *LCN2*) (Figures 2B-2I). In addition, ‘commensal’ *S. epidermidis* compared to strains more closely resembling pathogens (termed henceforth ‘inflammatory’ *S. epidermidis)* had notable differences in addition to increased activation of innate immunity M2 and M3 genes (Figure 2J); *IL23A*, a proinflammatory cytokine gene within the IL17 pathway was markedly and uniquely upregulated only by the latter. Together, these results demonstrate strong species and strain-level differences on RHE innate immunity.

**Figure 2.**
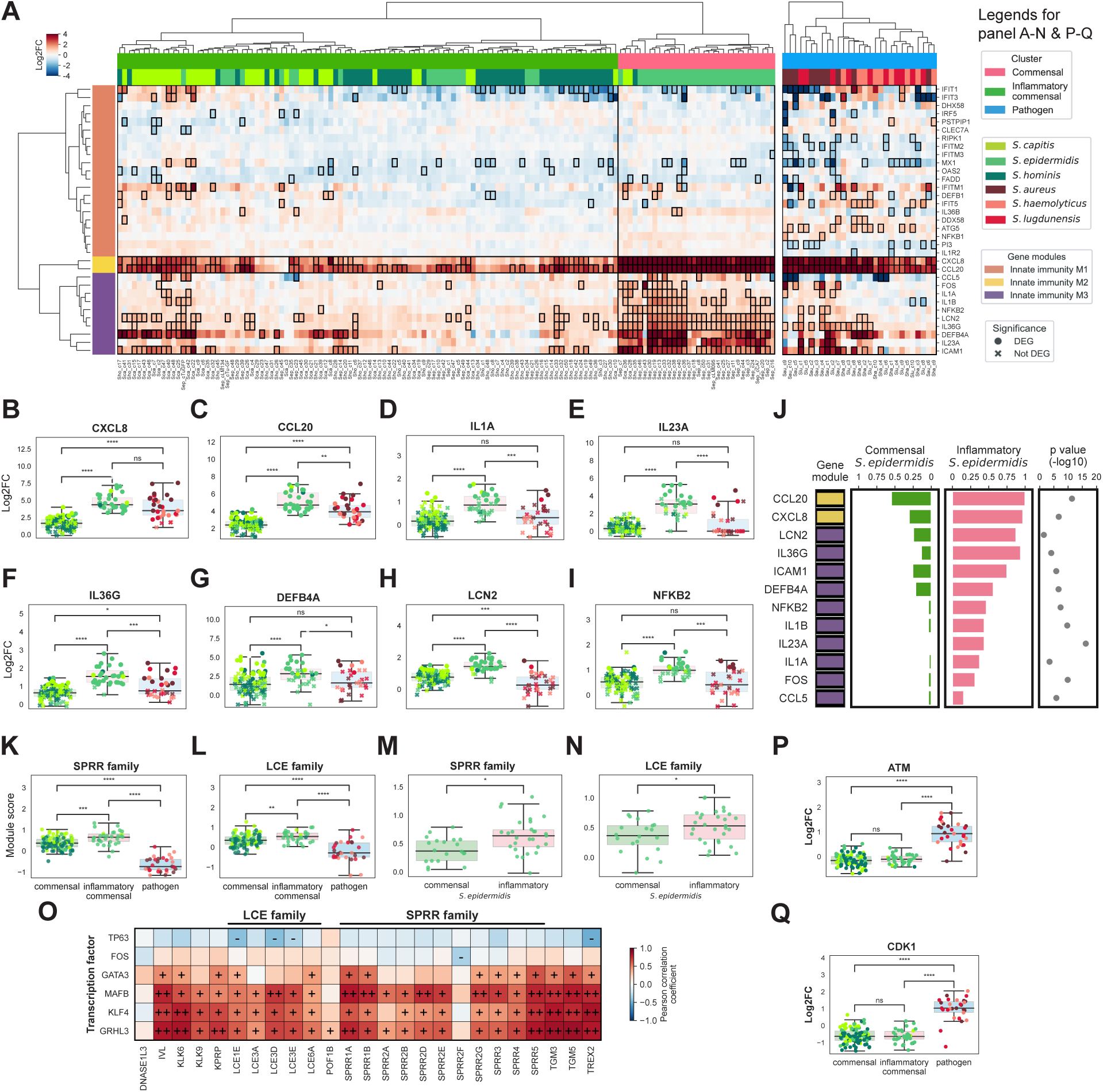
*Staphylococcus* species and strains drive differential responses in RHE for inflammatory, skin barrier, and proliferative programs. **(A)** Heatmap of log2FC transcriptional responses of a curated innate immunity gene panel in RHE following colonization by *Staphylococcus* strains (n=3 replicates per strain) relative to vehicle controls (n=14 replicates). Each column represents a different *Staphylococcus* strain treatment (colors, second row above heatmap), and each row corresponds to a gene. Outlined cells indicate statistical significance (FDR adjusted P-value ≤ 0.05 and |log2FC| ≥ 1). *Staphylococcus* strains were assigned into ‘commensal’, ‘inflammatory commensal’, or ‘pathogen’ clusters based on hierarchical clustering of their effects on RHE (colors in top row adjacent to heatmap). Similarly, innate immunity genes were clustered into ‘modules’ to reduce data dimensionality (colors in left column). For B-Q, legend panels are as in (A) and **** p ≤ 0.0001, *** 0.0001 < p ≤ 0.001, ** 0.001 < p ≤ 0.01, * 0.01 < p ≤ 0.05, ^ns^ p > 0.05. **(B-I)** Transcriptional responses (log2FC) of representative genes from innate immunity module M2 (B-C) and M3 (D-I), treatment vs. vehicle. Dots are colored by species as in (A); shape is statistical significance as in (A); grouping is by treatment cluster. **(J)** Innate immunity DEGs (FDR adjusted P-value ≤ 0.05 and log2FC ≥ 1) comparing commensal (n = 32) vs. inflammatory commensal (n = 16) *S. epidermidis* strains. Left column: colored by ‘gene module’ as in (A); central bar plots indicate the proportion of strain treatments within each cluster that induced upregulation of each gene in RHE; right plot shows P-value from Fisher’s exact tests. **(K-N)** Boxplots show transcriptional responses (module scores) of the SPRR and LCE gene families (L) in response to *Staphylococcus* strains (K, L, respectively) or *S. epidermidis* strains only (M, N, respectively). Module scores were calculated from the median of log2FC of genes in SPRR family (n=10) or LCE family (n=5). **(O)** Correlations between cornified envelope genes (x-axis) and reported transcriptional factors of cornified envelope genes (y-axis) in RHE colonized by *Staphylococcus* strains relative to vehicle controls. ++: Pearson correlation coefficient (r) ≥ 0.7, +: 0.3 ≤ r ≤ 0.7, -: −0.7 ≤ r ≤ −0.3. **(P-Q)** Boxplots showing log2FC of representative proliferative genes, *ATM* (P) and *CDK1* (Q). Boxes indicate the interquartile range (IQR, Q1 to Q3) of each group; center line indicates median.

#### Skin barrier effects

We similarly curated a panel of 311 skin barrier-related genes (Table S2) and created skin barrier modules to examine RHE’s barrier responses to colonization. Interestingly, species and strain-level clusters (Figure S2A) were largely conserved across skin barrier response genes. ‘Inflammatory commensals’ strongly upregulated skin barrier M1 and M4 genes, in contrast to the more moderate effect of ‘commensals’ and *S. epidermidis* strains clustering with ‘commensal’ species. These modules included multiple cornified envelope genes, primarily of the small proline-rich proteins (SPRR) and late cornified envelope (LCE) gene families (Figures 2K-2N), which are essential for cornified envelope structure and can function as antimicrobial proteins in skin barrier defense^31,32^. Furthermore, transcription factors (TFs) involved in regulation of cornified envelope genes, such as *GATA3*, *MAFB*, *KLF4*, and *GRHL3*, were strongly positively correlated with expression of cornified envelope genes, suggesting that some species- and strain-specific barrier effects may be mediated at least in part through these TFs (Figure 2O). Taken together, these results further support species- and strain-level differential effects on skin barrier processes in RHE.

#### Proliferative effects

We further examined ‘response-to-stimulus’ gene sets, which were uniquely modulated by ‘pathogenic’ *Staphylococcus* species but not commensals or inflammatory commensals (Figure 1C). Pathogens upregulated cell cycle regulator genes *ATM* and *CDK1* (Figures 2P-Q). *CASP4*, a proliferation-related gene reported to be downregulated in psoriasis^33^ was moderately downregulated by pathogens (Figure S2P). In contrast, *MKI67*, a canonical marker of keratinocyte proliferation^34^, was strongly induced by *S. aureus*, but not other pathogens or commensals (Figure S2Q). These findings underscore that pathogenic *Staphylococcus* preferentially induce proliferative programs in RHE, with *S. aureus* exerting particularly pronounced effects.

### Pathogenic versus non-pathogenic effects of *Staphylococcus* are recapitulated across RHE donors

Responsiveness to microbial colonization likely varies across keratinocyte donor, with likely age (neonatal vs. adult), skin site (foreskin vs. mature skin), and sex effects. To assess the generalizability of our findings, we constructed RHE from 3 additional donors: a neonatal foreskin (donor 827) and 2 adult female abdominal skin donors (27240 and 60780) (Figure S3A). Because the above variables are confounded in our model, we refer to them collectively as the ‘donor effect’. We selected a random set of 11 strains across 6 species for colonization (Table S3).

Overall, transcriptional responses were largely recapitulated across donors. Microbial effect held as the dominant influential factor driving expression of genes in innate immunity modules and SPRR and LCE gene families (5.0%-20.6% variance explained), consistent with a strong differential effect between commensal vs. pathogen categories. Donor was an influential factor (3.7%-8.5% variance explained) but exerted a much smaller influence (2.5%-3.7% variance explained) compared with the microbial effect (Figures 3A-3D). Altogether, approximately half of genes in the innate immunity and a quarter of genes in the skin barrier gene sets were shared across donors, with more genes shared between neonatal donors than adult donors (Figure S3B). Notably, differential responses to commensals and pathogens were largely conserved across both neonatal and adult donors (Figures 3E-3H and S3B), though interestingly, adult-derived RHE generally exhibited stronger inflammatory responses to colonization with larger effect sizes compared to neonatal donors (Figures 3I-J and S3B). Together, these results demonstrate that *Staphylococcus-*induced innate immune and barrier responses are largely robust across donors, with microbial identity as the primary determinant.

**Figure 3.**
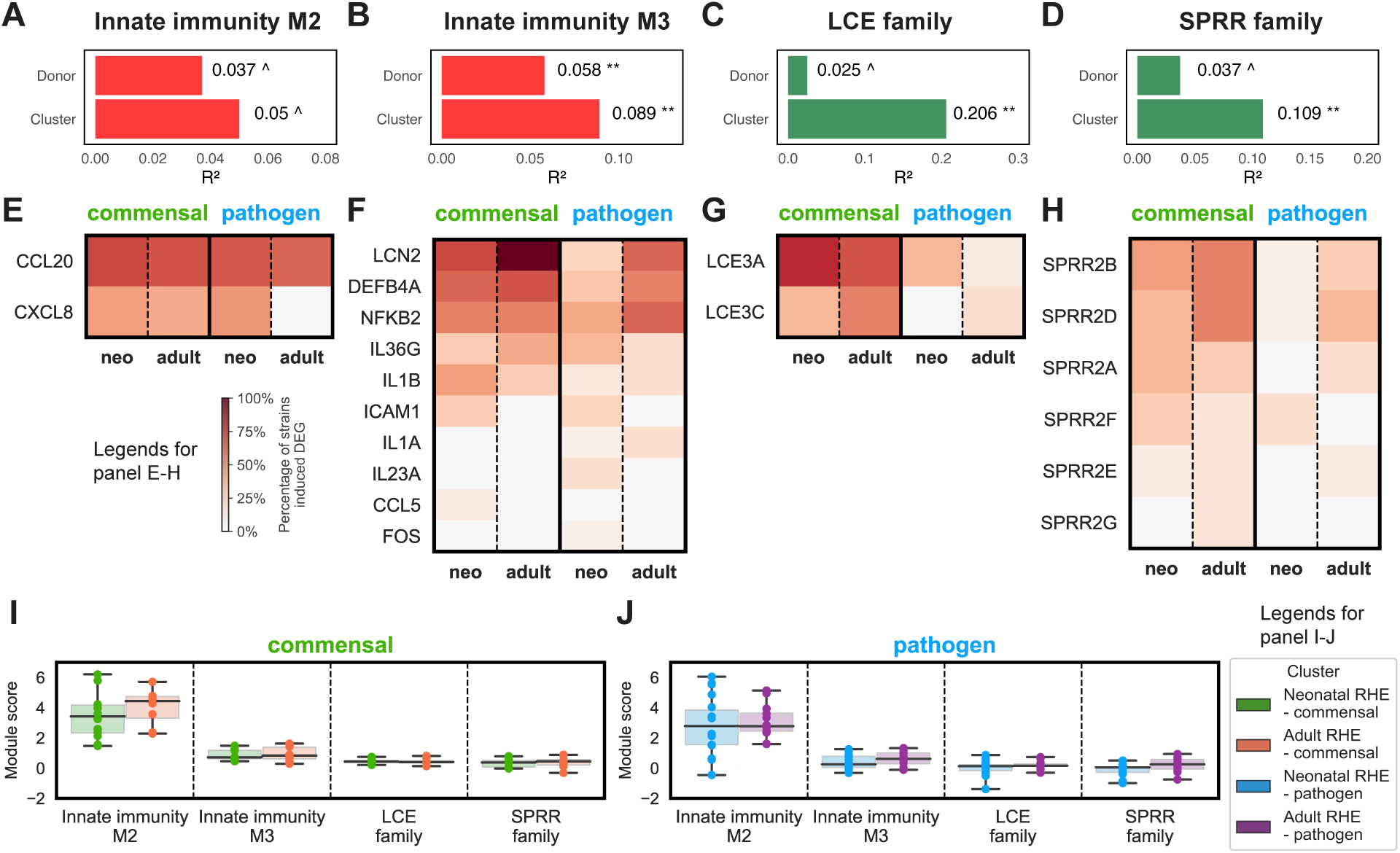
RHE response to *Staphylococcus* colonization is largely conserved across keratinocyte donors. RHE from neonatal (n=2) or adult (n=2) donors were colonized by a subset of 11 *Staphylococcus* strains from the original dataset (n=3 replicates per strain), or vehicle only. ‘Cluster’ reflects assignment from the original dataset. (A-D) Permutational multivariate analysis of variance (PERMANOVA) showing the proportion of variance (R^2^) in transcription of genes of innate immunity modules M2 (A), or M3 (B), or skin barrier LCE (C), or SPRR gene families (H) explained by *Staphylococcus* cluster and donor. Bars indicate R^2^ with associated P-values. **(E-H)** Heatmaps of proportion of *Staphylococcus* strains within ‘commensal’ vs. ‘pathogen’ clusters that significantly upregulated M2 (E), M3 (F), LCE (G) or SPRR gene families (H) in neonatal vs. adult RHE, to show consistency of response across donors. **(I-J)** Boxplots of module scores (median log2FC) of innate immunity modules and skin barrier gene families in neonatal (I) or adult (J) donors. IQR (boxes) and median (line) are shown. P-values: ** p ≤ 0.01, * 0.01 < p ≤ 0.05, ^ 0.05 < p ≤ 0.25.

### Gene-level, rather than global genomic or transcriptomic similarities, underlie differential effects on RHE

#### Global similarity

We next sought to identify staphylococcal factors underlying the differential RHE responses. First, to exclude differences in microbial growth rate (Figures S1B-S1D) as a factor, we modeled associations between RHE gene expression and bacterial load (CFUs at endpoint vs. inoculum). Few genes (23 for *S. aureus* and < 4 for all other species) correlated with CFUs out of 42,609 total human transcripts, none belonging to innate immunity or skin barrier gene panels (Figure S1E). This supports that RHE transcriptomic responses are largely independent of microbial growth. We then examined if differences in gene content were explanatory. Across strains, genome-wide gene presence-absence patterns were moderately correlated with RHE transcriptomes (r = 0.49, p < 1×10^-99^, Figure S4A), but notably, cluster differences accounted for greater divergence, rather than species or strain (Figures S4B-D and S4P-S4S). Surprisingly, phylogeny did not explain the stratification of *S. epidermidis* strains into inflammatory commensals and commensals (Figure S4E). Indeed, highly similar strains (r up to 0.91 Pearson correlation, representing as few as ∼100 genes differing), could elicit divergent responses whereas more divergent strains (r = 0.77, ∼400 genes differing) could produce similar phenotypes (Figure S4X). Thus, genome-wide differences only partially explained differential effects on RHE. Global transcriptomic similarity among strains was even less predictive of host responses (Figures S4F-J and S4T-U), suggesting that specific gene content differences rather than broad genomic or transcriptomic features might be a more important contributor to strain-specific effects on RHE.

#### Gene content differences

We next implemented a linear model to evaluate the degree to which specific gene contents at the species- or strain-level explained differential responses in RHE (Table S4). There were notable differences in KEGG modules, KEGG orthologs, virulence factors (VFs), and biosynthetic gene clusters (BGCs) between *Staphylococcus* in commensal vs. pathogen clusters (Figure 4A). As expected, pathogens were enriched for numerous virulence-associated features, such as the type VII secretion system, intercellular adhesion proteins, and capsular polysaccharides (Figures 4B-4C). Surprisingly, phenol-soluble modulins (PSMs) and lipases were more abundant in the commensal cluster, though these elements are typically virulence-associated (Figures 4D-4E). The staphyloferrin A BGC, which facilitates commensal colonization of bacteria on the skin^35,36^, was enriched in the commensal cluster (Figure 4F). Conversely, staphyloferrin B, a siderophore that promotes invasive bacterial growth^36^, was detected exclusively in certain pathogenic strains (Figure 4G).

**Figure 4.**
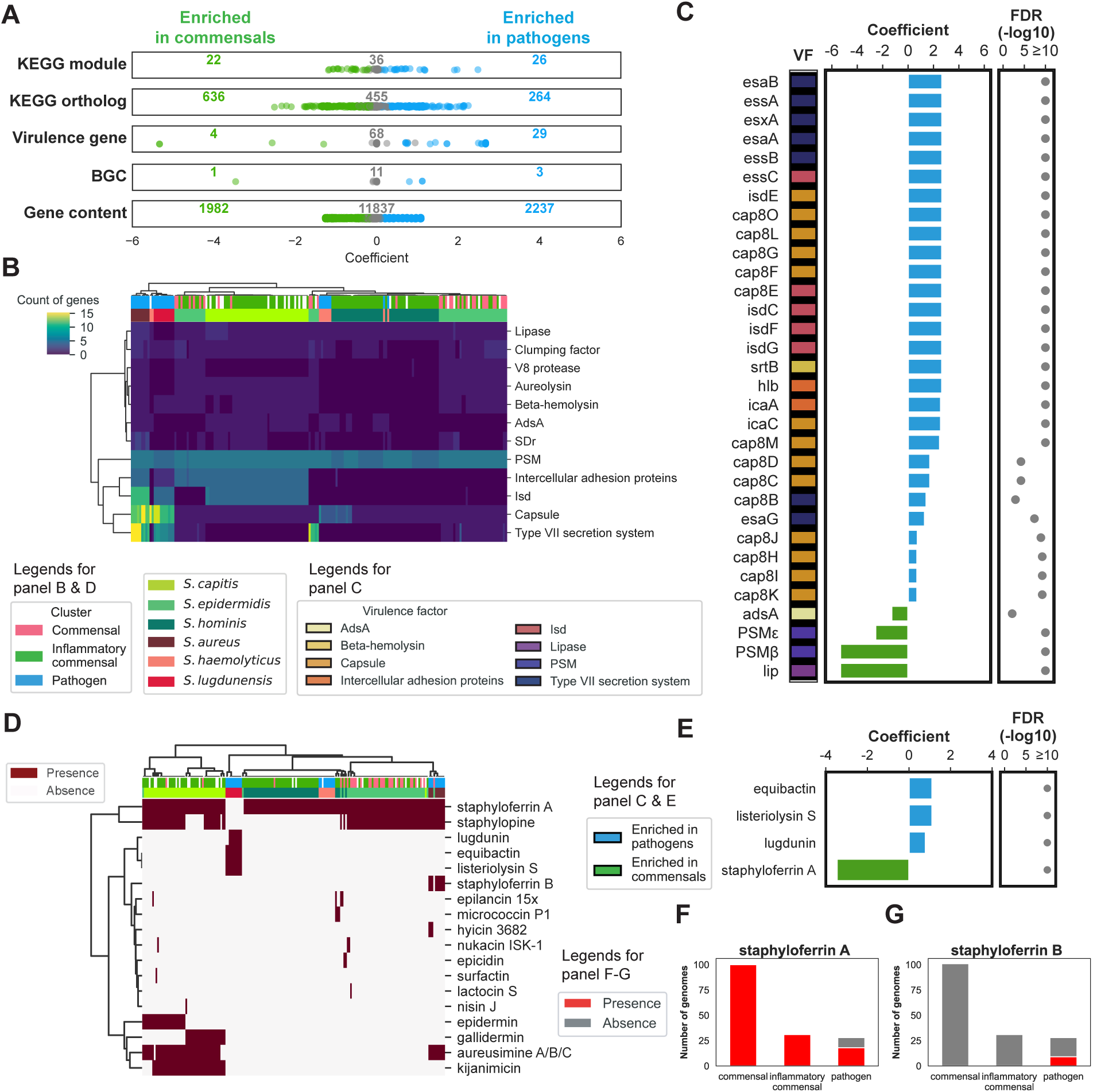
Genetic variation at the species level associated with *Staphylococcus* differential effects on RHE. **(A)** Linear regression analysis identifying *Staphylococcus* genomic features associated with commensal vs. pathogen clusters. Features included KEGG module completeness, presence/absence of KEGG orthologs (KOs), virulence factors (VFs), biosynthetic gene clusters (BGCs), or overall gene contents between *Staphylococcus* strains. Features were considered significantly enriched if FDR adjusted P-value ≤ 0.05. Points represent individual features, colored by enrichment; linear model coefficients reflect effect size with sign indicating enrichment in commensal (green) vs. pathogen (blue) clusters. Not significant (gray). Numbers indicating feature set size. **(B)** Heatmap shows strong segregation of commensal vs. pathogen clusters in prevalence of VFs (VFDB plus phenol soluble modulins), mapped from *Staphylococcus* genes across species and cluster. **(C)** Differential presence of VFs in commensal versus pathogen cluster. Center plot shows linear model coefficient (effect size) with sign (+/-) and colors showing enrichment in commensal vs. pathogen clusters. Left, VF functional category; right, FDR adjusted P. **(D)** Heatmap showing BGC prevalence in *Staphylococcus* genomes of the project annotated by strain and cluster. **(E)** As in (C), for BGCs, with **(F)** highlighting the prevalence of BGC staphyloferrin A and **(G)** B in *Staphylococcus* genomes.

At the strain level (Figure 5A), inflammatory *S. epidermidis* were enriched for genes associated with host interaction and stress adaptation, including *ricR* (copper resistance during infection^37^) and *essD* (ESS secretion system^38^) (Figure 5B). Notably, cell surface proteins *fbe* and *sdrF*, which mediate adhesion to keratinocytes^39–41^, were enriched in inflammatory strains and correlated with the expression of multiple proinflammatory genes in RHE (Figure 5C). For example, *IL23A*, which was uniquely upregulated by inflammatory *S. epidermidis*, was positively associated with the presence of both *fbe* and *sdrF* (Figure 5D).

**Figure 5.**
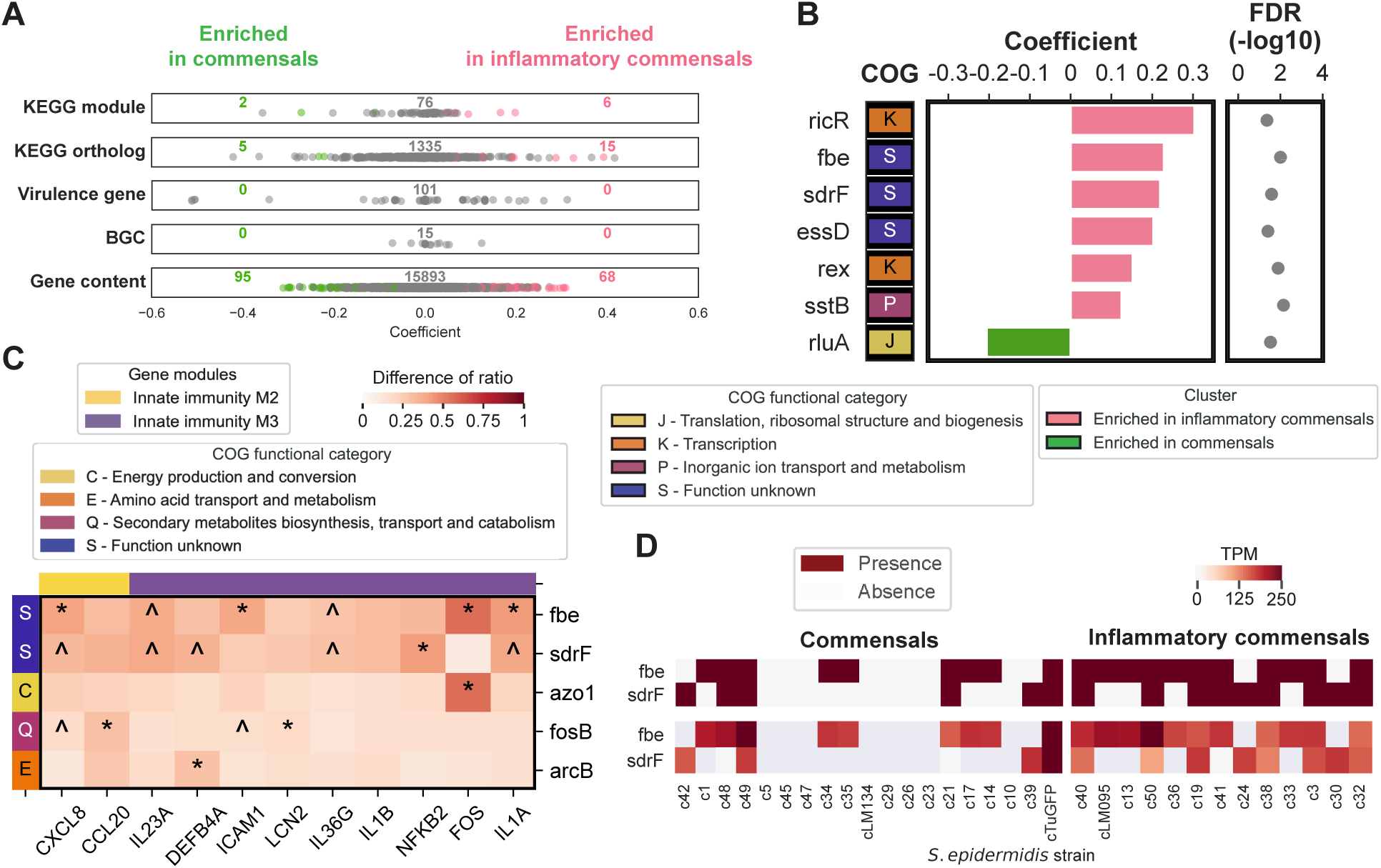
Genetic variation at the strain level associated with *S. epidermidis* differential effects on RHE. **(A)** Linear regression analysis identifying genomic features associated with commensal (green) vs. inflammatory commensal (pink) strain clusters, as in Figure. 4A. **(B)** Genes of differential presence between strains in commensal versus inflammatory commensal clusters. Center plot shows linear model coefficient (effect size) with sign (+/-) and colors showing enrichment in commensal vs. inflammatory commensal clusters. Right, FDR adjusted P; left; COG functional category. **(C)** Genes enriched in inflammatory *S. epidermidis* strains associate with upregulation of innate immunity genes in RHE. *S. epidermidis* gene presence was compared between strains that significantly upregulated innate immunity genes (DEG group) and those that did not (non-DEG group). For each innate immunity gene in Modules 2 and 3, associations with *S. epidermidis* genes were tested using Fisher’s exact test (2×2 contingency tables). Positively associated *S. epidermidis* genes (enriched in inflammatory strains) and their corresponding innate immunity genes are shown in the heatmap. Color indicates the difference of presence ratios (DEG − non-DEG). COG categories of *S. epidermidis* genes are annotated on the left, and innate immunity modules are indicated at the top. * p ≤ 0.05, ^ 0.05 < p ≤ 0.1. **(D)** Representative genes (*fbe* and *sdrF)* enriched in inflammatory *S. epidermidis* associated with *IL23A* induction in RHE. Top, gene presence of *fbe* and *sdrF* across *S. epidermidis* strains. Bottom, gene expression (Transcripts Per Kilobase Million (TPM)) of *fbe* and *sdrF*.

#### Transcriptional plasticity

To assess whether *Staphylococcus* transcriptional differences further contributed to host responses beyond gene content, we examined correlations between RHE transcriptomes and genes that were shared across strains but differentially expressed (prevalence >50% and actively transcribed in >80% of those genomes) (Figure S6F-I and Table S5). Genes involved in arginine metabolism (*arcC1* and *argF*), virulence (*bacA* and *capC*), and transcription (*hemA*, *hfq*, and *mfd*) had broadly positive correlations with innate immunity and skin barrier modules (Figure 6A). In contrast, *argG* (arginine metabolism) and *sufB* (iron uptake) were strongly negatively correlated with inflammatory genes such as *CXCL8* and *CCL20* (Figure 6B). Notably, these transcriptional patterns were specific to inflammatory commensals (Figures 6D-6H).

**Figure 6.**
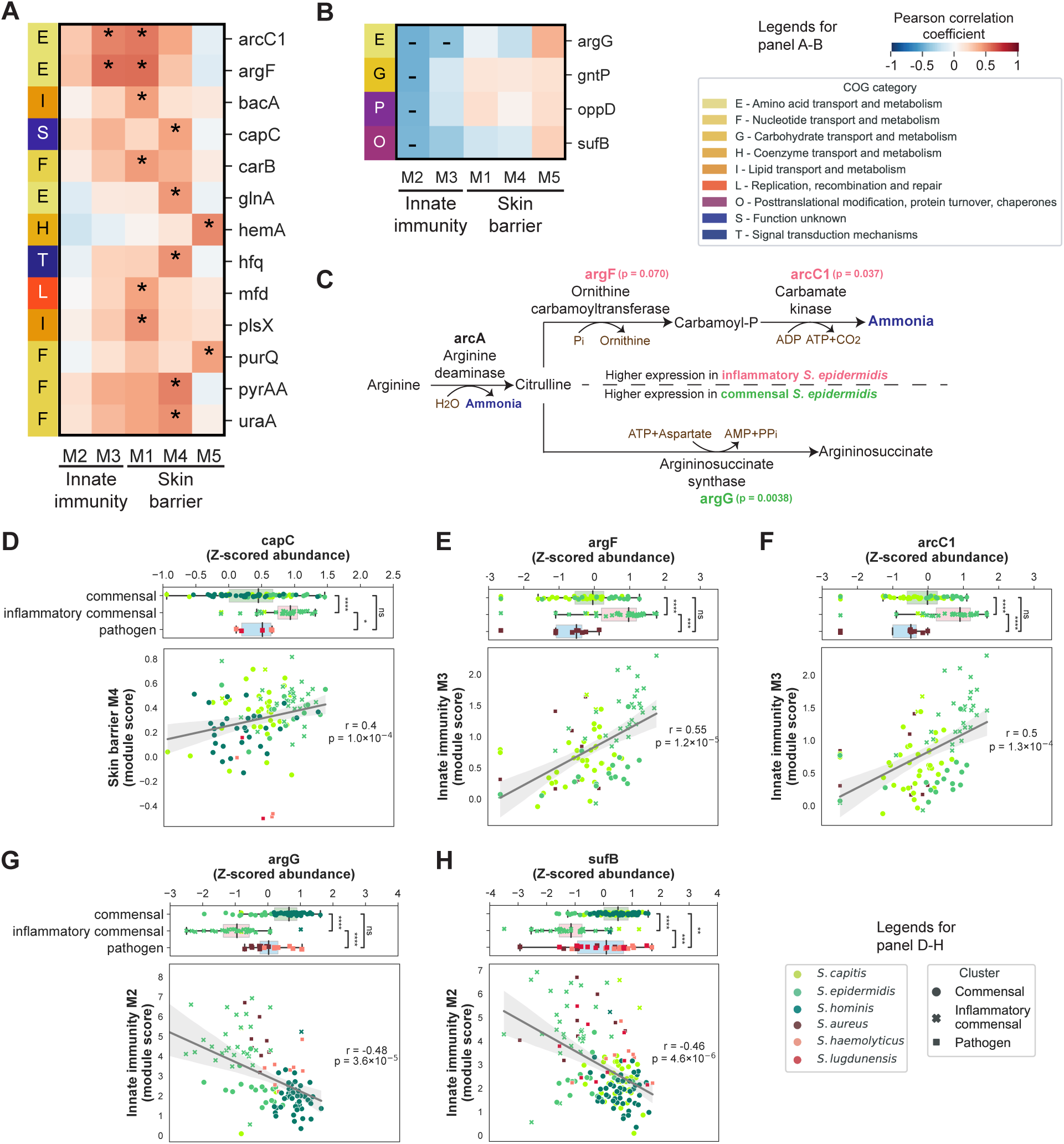
Species- and strain-transcriptional differences associate with their differential effects on RHE. (A-B) Co-expression between *Staphylococcus* genes (TPM followed by Z-score normalization, y-axis) and RHE innate immunity and skin barrier modules (median log2FC, x-axis), showing genes significantly (A) positively (*, Pearson correlation coefficient r ≥ 0.4, FDR adjusted P-value ≤ 0.05) or (B) negatively (-, r ≤ −0.4, FDR adjusted P-value ≤ 0.05). Left column, COG category of *Staphylococcus* genes. **(C)** Differential expression of the ammonia-producing arginine pathway between inflammatory and commensal *S. epidermidis* strains in RHE. Enzyme-coding genes are colored in red if significantly higher in inflammatory strains and green if higher in commensal strains (P-value ≤ 0.1). Pathway metabolites, reactions, and enzymes are shown. Genes in the ammonia-producing branch (*argF*, *arcC1*) are upregulated in inflammatory strains, whereas *argG* (ammonia-limiting branch) is higher in commensal strains. **(D-H)** Representative *Staphylococcus* genes significantly correlated with RHE transcriptional responses: (C) *capC* (skin barrier M4); (D) *argF* and (E) *arcC1* (innate immunity M3); (F) *argG*-and (G) *sufB* (innate immunity M2). Dot plots show relationship between *Staphylococcus* gene expression (TPM, x-axis) and RHE module activity (median log2FC, y-axis). Pearson correlation coefficient (r) and P-value (p) are indicated. Top box plots show gene expression stratified by cluster. Boxes represent IQR with median. **** p ≤ 0.0001, *** 0.0001 < p ≤ 0.001, ** 0.001 < p ≤ 0.01, * 0.01 < p ≤ 0.05., ^ns^ p > 0.05.

Focusing on *S. epidermidis* strain differences, strikingly, transcriptional differences in arginine metabolism distinguished inflammatory vs. commensal *S. epidermidis* (Figure 6C). Inflammatory strains had significantly elevated expression of *arcC1* and *argF*, which mediate conversion of arginine to ammonia. Commensal strains had higher expression of *argG*, which functions in the side branch of the pathway that consumes intermediates and limits ammonia production utilizing arginine. In contrast, genes involved in urea metabolism, an alternative pathway responsible for ammonia production and prevalent in our strains, exhibited neither strong correlations nor differential expression between *S. epidermidis* strains (Figure S6L and S6N). Collectively, these results identify discrete mechanisms and transcriptional programs that underlie strain-specific inflammatory potential, highlighting differential regulation of arginine metabolism to favor ammonia-producing pathways in inflammatory strains, and distinct repertoires of virulence and adhesion factors as important factors driving differential *Staphylococcus-*RHE interactions.

### *Staphylococcus* strain-level metabolic and phenotypic variation underlies differential effects on RHE

To functionally validate our multi-omics predictions, we assessed strain-level phenotypes and metabolic outputs associated with the inflammatory effects of *S. epidermidis* on RHE (Table S7). Arginine metabolism has been implicated in *S. epidermidis* biofilm formation^42^, which is strongly associated with pathogenicity, and thus arginine availability might be an environmental cue modulating *S. epidermidis’* inflammatory potential. We investigated whether inflammatory commensal *S. epidermidis* exhibited greater virulence-associated phenotypes than commensal strains. Indeed, approximately half of the inflammatory commensals formed biofilms, compared to only a quarter of commensal strains (Figures 7A and S7A). In a bloodstream infection model^43^, inflammatory commensals also exhibited robust survival similar to the canonical pathogen *S. aureus* USA300LAC^44^, with CFU levels at 24 hours comparable to the initial inoculum after a transient early decline (Figures 7B–C). In contrast, commensal *S. epidermidis* strains failed to recover after the initial reduction, an effect that was rescued under complement-depleted conditions (Figures 7D–E). Together, these phenotypic differences support our classification of *S. epidermidis* strains into inflammatory and commensal groups.

**Figure 7.**
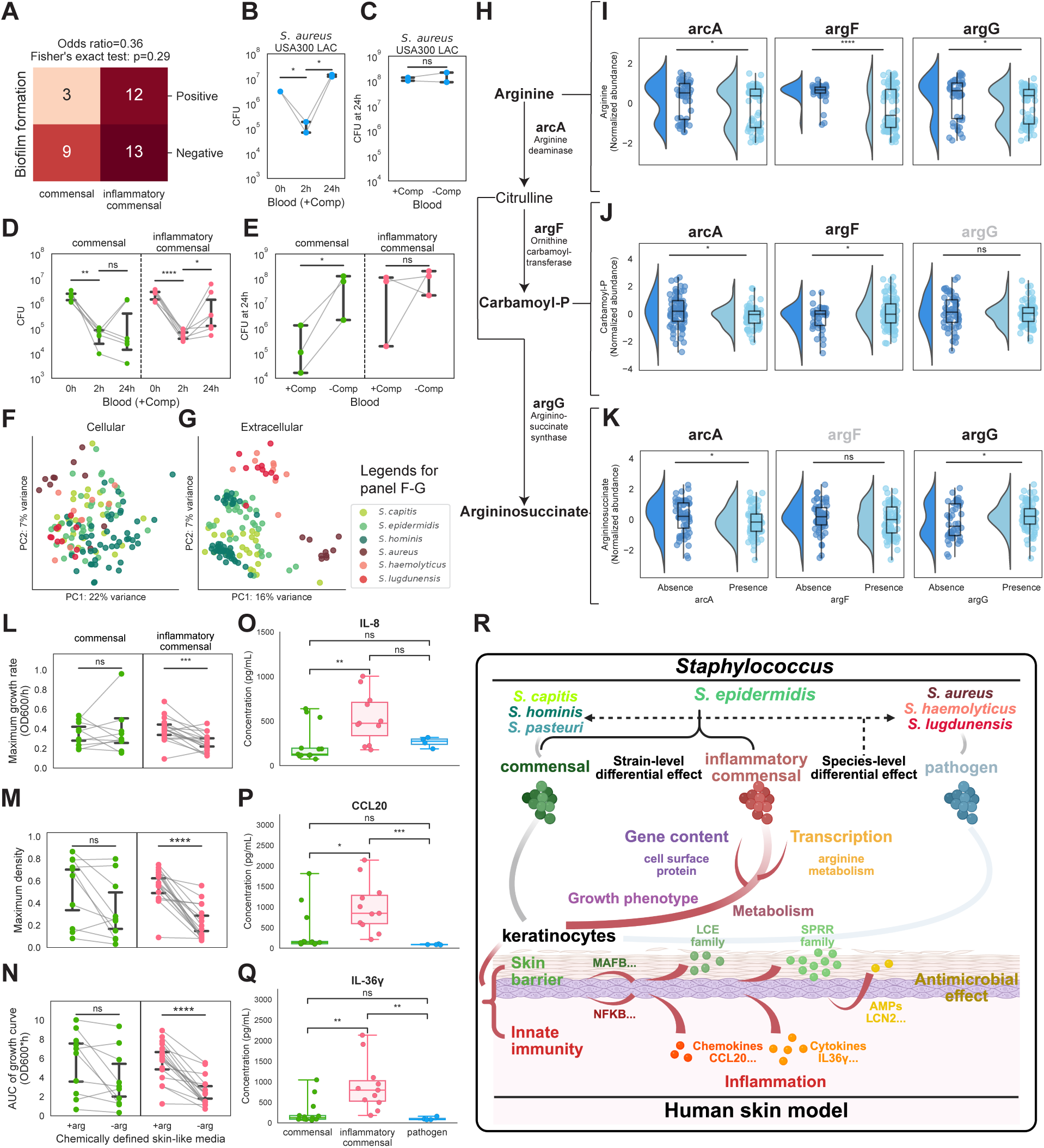
*Staphylococcus* strain-level metabolic and phenotypic differences underlie their differential effects on RHE. **(A)** Biofilm formation ability of commensal vs. inflammatory *S. epidermidis* strains. Biofilm positive strains were defined relative to a canonical biofilm-forming strain (Sep_1457). Numbers indicate strain counts per category and P-value and odds ratio calculated using Fisher’s exact test. **(B-E)** Survival ability in blood as measured by CFUs at 0, 2, and 24h of *S. epidermidis* strains (D) with respect to a bloodstream infection *S. aureus* strain (USA300LAC, (B)) with or without complement (+/-Comp) depletion (C, E). Points represent individual strains (or replicates for *S. aureus)* and boxes indicate IQR with median. Lines connect matched conditions. Significance was assessed by paired t-test with Bonferroni correction. **(F-G)** PCA plot of *Staphylococcus* cellular (F) or extracellular (G) metabolomes. *Staphylococcus* metabolomes were inferred by metabolic feature intensity within untargeted metabolomes. **(H)** Diagram of arginine metabolism pathway highlighting reactions catalyzed by *arcA*, *argF*, and *argG*. **(I-K)** Violin plots showing the metabolite abundance of arginine (I), carbamoyl-P (J), and argininosuccinate (K) in strains with or without *arcA*, *argF*, or *argG.* Points indicate normalized metabolite intensity; violins show distributions with embedded boxplots (median and IQR). Gene labels are black if acting on or downstream of the metabolite, grey otherwise. **(L-N)** Growth dynamics of inflammatory vs. commensal *S. epidermidis* strains in skin-like media with (+Arg) or without (-Arg) arginine. Maximum growth rate (L), maximum density (M), and area under the curve (AUC) (N) are derived from OD600 measurements every 0.5h over 10 hours at 37C. Points represent strains with paired conditions connected with lines. Statistical significance was determined by paired *t*-test followed by Bonferroni correction. **(O-Q)** Protein expression of cytokines IL-8 (O), CCL20 (P) and IL-36γ (Q) in the basal media of *Staphylococcus*-colonized RHE was measured by ELISA (n=3-4 replicates per strain in an independent experiment colonizing 3 inflammatory and 4 commensal *S. epidermidis* strains and *S. aureus* as a reference). Points represent individual replicates colored by cluster and boxes show IQR with median. **(R)** Model summarizing species- and strain-level effects of *Staphylococcus* on RHE. Species-level differences distinguish commensal and pathogenic effects, while strain-level variation within *S. epidermidis* identifies inflammatory strains that activate a proinflammatory transcriptional program including upstream regulators (e.g. NFKB), chemokines (e.g., CCL20), cytokines (e.g., IL36γ) and downstream antimicrobial effectors (e.g. LCN2). Inflammatory *S. epidermidis* also enhances skin barrier defensive genes, particularly members of the LCE and SPRR gene families, which contribute to both barrier integrity and antimicrobial defense, and are regulated by transcription factors such as MAFB. These effects are linked to differences in gene content (e.g., cell surface genes), transcriptional programs, metabolism (e.g., arginine pathway), and pathogen-like growth phenotypes. P-values: **** p ≤ 0.0001, *** 0.0001 < p ≤ 0.001, ** 0.001 < p ≤ 0.01, * 0.01 < p ≤ 0.05., ^ns^ p > 0.05.

To identify potential metabolic drivers, we performed untargeted metabolomics on isolates grown in rich media to avoid complicating host metabolites. Metabolite profiles varied strikingly across species and strains, with extracellular metabolites (supernatant) showing clear segregation by cluster classification and intracellular metabolites (pellet) showing greater overlap between species (Figures 7F-7G). Interestingly, pathogens exhibited a similar dynamic range in metabolomic variability as commensals and inflammatory commensals despite being represented by fewer strains (Figures S4V-S4W). Surprisingly, genetic similarity was only moderately positively correlated with metabolic similarity (Figure S4K), indicating that gene content differences alone do not fully explain metabolic output.

Given the importance of arginine metabolism in our transcriptional analyses, we next assessed this pathway in *S. epidermidis* strains (Figure 7H). Strains harboring *arcA* showed decreased arginine abundance, consistent with its role in converting arginine to citrulline and ammonia (Figure 7I). Presence of *arcA* did not influence the abundance of carbamoyl-P and argininosuccinate, two downstream metabolites in the pathway (Figures 7J-K). In contrast, *argF* was correlated with higher carbamoyl-P and lower arginine levels, consistent with its role in converting citrulline to carbamoyl-P (Figure 7J). Similarly, *argG* presence correlated with higher argininosuccinate and lower arginine levels, consistent with its role in converting citrulline to argininosuccinate (Figure 7K). Functionally, inflammatory and commensal *S. epidermidis* grew comparably in chemically defined skin-like media^45^ replete with arginine. However, under arginine limitation, inflammatory *S. epidermidis* exhibited significantly impaired growth (maximum growth rate, maximum density, and area under the growth curve), whereas commensal *S. epidermidis* showed no significant impairment (Figures 7L–7N). These findings demonstrate that strain-specific metabolic programs, particularly differential utilization of arginine, contribute to phenotypic variation and inflammatory potential in *S. epidermidis*.

### Validation cohorts support strain-level differential effects of *Staphylococcus* on RHE

Finally, to assess the robustness and generalizability of our findings, we evaluated independent validation cohorts across experimental and biological conditions. First, to account for variability in RHE differentiation across batches, we repeated the colonization experiments with a random subset of 9 strains from 3 commensal species (Figure S7D). A classification model trained on the original dataset (see Methods) achieved 94% accuracy in predicting strain effects in this independent cohorts (Figure S7E), correctly recapitulating inflammatory commensal and commensal assignments (Figure S7F). Furthermore, the expression patterns of innate immunity modules (M2 and M3) and skin barrier modules (M1 and M4) were consistent between cohorts (Figures S7G-S7L). Second, to test generalizability to previously unobserved strains, we profiled RHE responses to colonization by 7 additional strains of *S. pasteuri*, another hypothesized commensal species (Figure S7D). All 7 strains were classified as commensals (Figure S7F) with transcriptional effects on innate immunity M2 and M3 resembling other commensals (Figures S7M-S7N), supporting the versatility and reproducibility of our dataset.

We then validated functional outputs by measuring cytokine secretion following colonization with a third independent representative set of strains from the different clusters (Figure S7O). Consistent with our transcriptomic data, IL-8 protein levels were significantly higher in samples treated with pathogens and inflammatory *S. epidermidis* than in those treated with commensal *S. epidermidis* (Figure 7O). In contrast, chemokine CCL20 and cytokine IL-36γ were selectively increased in inflammatory *S. epidermidis* relative to both commensal *S. epidermidis* and pathogenic strains (Figures 7P-7Q). Collectively, these results underscore that strain-level differences in *Staphylococcus* elicit reproducible and generalizable effects on RHE across experimental conditions, extending from transcriptional programs to functional protein outputs and reinforcing their impact on skin inflammatory and barrier programs on human skin (Figure 7R).

## DISCUSSION

Human skin harbors diverse *Staphylococcus* species, each with extensive strain variation. Because of this remarkable scope, systematic investigation of how these differences influence skin biology has been limited. Here, we present a large-scale, multi-‘omics investigation of strain-level interactions between numerous *Staphylococcus* species and human skin, revealing substantial functional heterogeneity across both species and strains.

A central finding of this study is that skin responses are strongly shaped at both at the species and strain level. Species such as *S. lugdunensis, S. haemolyticus,* and canonical pathogen *S. aureus* elicited a stronger proinflammatory response in the skin model than typical commensal species including *S. hominis, S. capitis, and S. pasteuri*. However, a subset of *S. epidermidis* strains induced transcriptional programs more closely resembling those of pathogens, which we termed ‘inflammatory commensals’. These responses included activation of a proinflammatory gene network with key regulators (*NFKB2* and *FOS*), cytokines (*IL1A* and *IL36G*), chemokines (*CXCL8* and *CCL20*), and antimicrobial peptides (*DEFB4A* and *LCN2*) typically associated with skin infections and inflammatory skin diseases^46–48^. Notably, *IL23A*, a member of the IL17 gene family^49^, was uniquely upregulated by inflammatory *S. epidermidis*, paralleling findings in *S. aureus*-induced skin infections and psoriasis-like symptoms^50,51^. However, only trace IL-23 protein was detected in basal media of *Staphylococcus*-colonized RHE, likely due limited expression of the shared *IL12B* subunit, indicating that further model optimization may be necessary to recapitulate disease-relevant IL-23 responses. In the context of skin disease, the increased secretion of CXCL8, CCL20, and IL-36γ in response to colonization by inflammatory commensals may orchestrate downstream tissue resident immune activation, including neutrophils^52^ and Th17^53^ cells.

In parallel, we observed coordinated modulation of skin barrier programs. Commensal species broadly upregulated barrier associated genes whereas pathogenic species elicited minimal effect. In contrast, inflammatory *S. epidermidis* upregulated cornified envelope gene families, including LCE^54^ and SPRR^31,55^, which have both structural and antimicrobial functions. This response was associated with transcription factors such as *GATA3*, *MAFB*, *KLF4* and *GRHL3*^56,57^. Notably, *GATA3* regulates cornified envelope genes downstream of TLR2 activation^58^, a key keratinocyte receptor for staphylococcal ligands such as lipoprotein^59^, suggesting a link between microbial sensing and barrier activation. These findings support a model in which keratinocytes mount a coordinated inflammatory, antimicrobial, and barrier responses that vary based on the functional properties of the colonizing strains, with potential consequences for skin homeostasis and microbial ecology.

Our extensive comparative genomics analyses identified multiple gene content differences that may underlie these phenotypes. Pathogenic strains were enriched for expected features such as virulence factors and biosynthetic gene clusters facilitating skin colonization^35,36^. *S. epidermidis* and *S. capitis* were enriched in more virulence-associated processes compared to *S. hominis*, including the VraFG transporter, ABC transport system permeases, and multidrug resistance protein MFS transporter (Figures S5I-S5J). However, despite these similarities, *S. capitis’* effects on skin were more similar to *S. hominis*, suggesting that the inflammatory commensals might possess unique combinations of features that drive their effects. Indeed, inflammatory *S. epidermidis* uniquely possessed a siderophore associated with invasive bacterial growth^35,36^, *essD*, a component of the ESS secretion system linked to virulence^38^, and *fbe* and *sdrF* surface protein genes that facilitate *Staphylococcus* adhesion to keratinocytes^41,60^. These findings support that commensal *Staphylococcus* species can serve as reservoirs for pathogenicity-associated genes^61–65^, while highlighting that specific gene repertoires rather than overall virulence gene enrichment can determine strain-level effects on skin.

Our dual transcriptomic analysis further revealed that variation in host responses was driven by differential expression of specific staphylococcal genes and not necessarily global transcriptional activities. For example, virulence factors *capC*^66^ and *bacA*^67,68^ were upregulated in inflammatory commensals and positively correlated with RHE’s inflammatory and skin barrier gene expression. Strikingly, inflammatory and commensal *S. epidermidis* strains differed in their regulation of arginine metabolism. Inflammatory strains preferentially expressed pathways purported to promote ammonia production, whereas commensal strains favored pathways that would divert intermediates and limit ammonia production, transcriptional differences supported by metabolomic and functional assays. Arginine metabolism has been implicated in microbial pathogenicity through its capacity to generate ammonia^50,69^, and ammonia-related pathways were enriched in the skin microbiota of atopic dermatitis patients, potentially contributing to elevated pH^70,71^. Given this potential role of arginine metabolism as a driver of *S. epidermidis’* inflammatory potential on skin, modulating arginine metabolism or availability in the skin could be a strategy to selectively limit the effects of inflammatory strains while supporting beneficial commensals.

In conclusion, our study establishes a multi-omics framework for understanding how strain variation across *Staphylococcus* species shapes skin biology and provides new insights into the continuum of commensalism and pathogenicity of these keystone skin microbiota. By integrating genomic, transcriptional, metabolic, and phenotypic data, we show that host–microbe interactions on the skin are shaped by not only species but also discrete strain-specific properties. Our findings underscore the importance of strain-level analyses, provide insights into mechanisms of microbe-skin interactions, and offer a robust platform to further dissect the role of the skin microbiome in skin health and disease.

### Limitations of the study

We note several limitations of this study. First, we selected a single treatment timepoint—18 hours—for the primary screening dataset and all subsequent validation experiments. This choice was guided by a preliminary time series experiment, which identified 18 hours as an intermediate inflammatory response approximating early colonization. However, a more extensive time series would be valuable to capture temporal dynamics of host-microbe interactions. Second, while we utilized RHE as a model system that recapitulates key features of the epidermal barrier, it lacks the full complexity of human skin. More advanced systems incorporating dermal layers, immune cells, and skin appendages would better capture multicellular interactions and immune responses to microbial colonization. Third, our use of bulk transcriptomics was appropriate given the relatively homogeneous RHE model. Future studies incorporating spatially resolved approaches, such as spatial transcriptomics, would be valuable for resolving layer-specific responses and cell-cell communication in more complex skin models. Fourth, our metabolomics analyses were performed on *Staphylococcus* cultured in rich media rather than in co-culture with RHE, reflecting challenges in differentiating microbial from host-derived metabolites in untargeted datasets. Targeted metabolomics approaches and improved annotation will be important for accurately defining microbial metabolic activity in the skin microenvironment, which are likely highly sensitive to environmental cues. Finally, our experiments examine individual strains in isolation and do not capture the ecological interactions inherent to complex skin microbiota. In vivo, inter-strain, and interspecies interactions may influence colonization dynamics, metabolic outputs, and host responses. Incorporating community-level models is an important next step to understand how these strain-specific effects manifest in the context of the native skin microbiome.

## METHODS

### Generation of skin-derived *Staphylococcus* strain collection

A collection of 187 skin-derived isolates of seven *Staphylococcus* species was selected from a biorepository of 2,812 whole genome sequenced strains isolated from healthy human skin, patients with congenital ichthyosis, and miscellaneous other human skin sources^28–30^. Genomes of these strains were placed on a phylogenetic tree built on gene content differences and ‘cut’ (cuttree option in R) to identify representative isolates from different genetic lineages to maximize genetic diversity. It includes 50 * *S. epidermidis* strains (Sep_c1-Sep_c50), 50 * *S. capitis* strains (Sca_c1-Sca_c50), 50 * *S. hominis* strains (Sho_c1-Sho_c50), 7 * *S. pasteuri* strains, 10 * *S. aureus* strains (Sau_c1-Sau_c10), 10 * *S. haemolyticus* strains (Sha_c1-Sha_c10), 10 * *S. lugdunensis* strains (Slu_c1-Slu-c10). Additional 4 *S. epidermidis* lab strains (*S. epidermidis* Tü3298-GFP (Sep_cTuGFP), *S. epidermidis* CA7 (Sep_cCA7), *S. epidermidis* NIHLM095 (Sep_cLM095), and *S. epidermidis* NIHLM134 (Sep_cLM134)) were further included.

### Reconstructed human epidermis (RHE) cell culture cultivation

9mm primary normal human RHE tissue cultures (EpiDerm) were obtained from MatTek Corporation. All batches were grown using cells from EpiDerm standard donor, a healthy neonatal male, except for the alternative donors used for examining donor-specific responses, which included an additional neonatal male and two adult female donors. RHE cultures were cultured according to the manufacturer’s directions. Briefly, upon arrival, the RHE cultures were placed in 6-well plates with 1mL of warmed antibiotic-free EpiDerm Maintenance Media or the equivalent EpiDerm Assay Media basally per well. The basal media was replaced daily, and the RHE cultures were kept at 37°C with 5% CO_2_.

### Colonization of RHE by *Staphylococcus* isolates

#### 187-isolate experiment

For each microbial isolate, a single colony was grown overnight in sterile 1X Tryptic Soy Broth (TSB, Becton, Dickinson and Company; #211825). 10^7^ colony-forming units (CFU) were taken from each liquid culture and washed with ultrapure water (Fisher Scientific; #AAJ71786AP), with a final concentration of 10^7^ colony-forming units in 120µL. The RHE cultures were then dosed with 120µL of microbial isolate/vehicle. Dosed RHE cultures were incubated for 1 hour at 37°C, then inoculum was removed to restore the air liquid interface. RHE were incubated with remaining bacteria for 18 hours prior to harvest. Excess inoculum or vehicle controls were serially diluted in sterile phosphate-buffered saline (PBS) and grown on Trypticase Soy Agar (TSA) with 5% sheep’s blood (Fisher Scientific; #221261) to determine input CFUs. At harvest, 200µL of PBS was added to the apical surface of each RHE, pipette-mixed, then removed. This PBS wash was serially diluted and plated on TSA with 5% sheep’s blood or TSB supplemented with Bacto Dehydrated Agar (Fisher Scientific; #214010) for CFU counts. The central region of each RHE was then excised using a 6mm biopsy punch (Integra, #33-26) and transferred into 140 µL of Buffer RLT (Qiagen; #79216) supplemented with 1% β-mercaptoethanol for RNA preservation. The Buffer RLT-tissue solution was frozen at −80°C until RNA extraction. *S*. *epidermidis* Tü3298-GFP colonized RHE were visualized under blue light to confirm colonization. *Donor variation experiment:* A set of 11 strains of 6 species of *Staphylococcus* (2 * *S. epidermidis* strains, 1 * *S. capitis* strain, 2 * *S. hominis* strains, 2 * *S. aureus* strains, 2 * *S. haemolyticus* strains and 2 * *S. lugdunensis* strains) from the 180-strain collection were picked randomly. As above, each strain was applied to RHE derived from 4 different donors (standard donor (neonatal male), donor 827 (neonatal male), donor 27240 (35-year-old female), and donor 60780 (45-year-old female)) (MatTek Corporation) in biological quadruplicate.

#### Validation experiment 1

A set of 9 strains from 3 species of *Staphylococcus* (4 * *S. epidermidis* strains, 2 * *S. capitis* strains and 3 * *S. hominis* strains) were randomly selected from the set of 180, plus 7 *S. pasteuri* strains. Each strain was applied to RHE in biological quadruplicate and treated and processed for CFU analysis and RNA-seq as above.

#### Validation experiment 2 and cytokine analysis

A set of 8 strains including 3 *S. epidermidis* strains from the inflammatory commensal cluster, 4 *S. epidermidis* strains from the commensal cluster and 1 *S. aureus* strain from the pathogen cluster, were selected for RHE colonization as above. In addition to CFU analysis and RNA-seq, basal media from each RHE was frozen at −80°C for cytokine analysis. ELISA was performed with DuoSet ELISA Kits (R&D Systems; #DY360, #DY206, #DY2320-05 and #DY1290) according to the manufacturer’s recommendations for the presence of CCL20, IL-8, IL-36G and IL-23.

### RNA extraction, library preparation, and RNA-seq

All RNA extraction and sequencing library preparation steps were performed in a sterile tissue culture hood. Prior to extraction, samples were mechanically homogenized to ensure efficient disruption of the epidermal tissue; tissues were beaten in a Qiagen TissueLyser for 6 minutes at 30 hZ with 100uL of 0.1mm glass beads (Biospec Products cat 11079101) in ∼140uL of RLT+1%BME (Qiagen). RNA was then extracted using the RNeasy 96 QIAcube HT kit (Qiagen, #74171) according to the manufacturer’s directions. Samples were eluted in nuclease-free water (Qiagen, #129115) and frozen at −80°C until sequencing preparation. RNA quality was evaluated using the 4200 TapeStation System (Agilent Technologies) with the High Sensitivity RNA ScreenTape Analysis or RNA ScreenTape Analysis. RINs ranged from 1.7-9.9 with an average of 8.4 and a median of 8.9. RNA quantity was measured on the Qubit 2.0 Fluorometer (Thermo Fisher Scientific). The sequencing libraries were prepared with NEBNext rRNA Depletion Kit v2 (New England Biolabs; #E7400X) and NEBNext Ultra II Directional RNA Library Prep Kit for Illumina (New England Biolabs; #E7760L) following the manufacturer’s directions. Library quality was evaluated using the 4200 TapeStation System with the High Sensitivity D1000 ScreenTape Assay. Library quantity was measured on the Qubit 2.0 Fluorometer. Samples were sequenced using Illumina NovaSeq. Reads per sample ranged from 459K to 152 million with an average of 32 million reads and median of 32 million reads.

### Transcriptional profiling of RHE transcriptome

RNA-seq reads were processed with trimmomatic 0.39^72^ to remove low-quality reads. Quality-controlled reads were mapped to CHM13v2.0 reference genome with STAR 2.7.1a^73^ to retrieve human reads. Read counts aligned to each gene was computed using featureCounts v1.6.4^74^ (from Subread1.6.4). Genes with read counts lower than 10 in more than 50% samples (including both strain-treated RHE and vehicle controls) were filtered out prior to differential expression analysis. Potential batch variation was removed using RUVg (from RUVseq 1.38.0^75^) with the top 5000 stably expressed genes for each comparison. Differential gene expression was analyzed using DESeq2 1.44.0^76^. Log2 fold change (FC) for each gene were estimated using DESeq2’s normalized read counts. Genes with |log2FC| ≥ 1 and FDR ≤ 0.05 were considered differentially expressed.

### Gene Ontology enrichment analysis

Gene Ontology(GO) enrichment analysis was performed on upregulated or downregulated DEGs calculated by DESeq2^76^ using enrichGO (from clusterProfiler 4.12.0^77^). Gene sets of FDR ≤ 0.05 were determined as significantly differentially expressed gene sets.

### Curated gene panels for RHE RNA-seq data

The innate immunity gene panel was adapted from MSigDB 2025.1.Hs^78^ with gene ontologies (GO:0005125, GO: 0061844, GO: 0019730, GO: 0071347, GO:1990266). The skin barrier gene panel was adapted from the literature^31,32,79^.

### Microbial population (CFU) confounder analyses

CFU confounder analysis was performed using MaAsLin2 1.18.0^80^. RHE transcriptomes were analyzed using MaAsLin2, with CFU counts of microbial treatments at the end point (18 hour) and species included as fixed effects and batch included as a random effect. Genes were considered as confounded by CFU if counts of genes were significantly associated with CFU counts within the dataset.

### Clustering analysis for dimensionality reduction of RHE innate immunity and skin barrier transcriptional responses

Responsive genes within innate immunity and skin barrier gene panels were determined as expressed in ≥ 50% of treatment comparisons and were significantly differential expressed in ≥ 2 comparisons. The log2FC matrix of responsive gene lists to strain-to-vehicle comparisons was further processed by hierarchical clustering on both axes using an implanted method in seaborn^81^ Python package with method = ward. The clusters of strain-to-vehicle comparisons were defined as commensal and inflammatory commensal. The clusters of genes were defined as innate immunity modules (innate immunity M1-M5) for the innate immunity gene panel and skin barrier modules (skin barrier M1-5) for skin barrier gene panel.

### Whole genome sequencing, assembly, and annotation

Whole-genome sequencing of *Staphylococcus* strains was performed using a paired-end 2 × 150 bp protocol on the Illumina NovaSeq 6000 platform, as previously described^16^. Low quality bases and sequencing adapters were removed from the sequencing reads from whole genome sequencing using PRINSEQ-lite 0.20.4^82^ and trimmomatic 0.39^72^. Then filtered sequencing reads were used to assembled using SPAdes 3.7.1^83^ with default parameters. Contig quality was assessed using QUAST^84^.

Gene coding sequences were first predicted from the assembled genomes using Prokka 1.14.6^85^ with kingdom = Bacteria and genus = Staphylococcus. The pan- and core-genomes were further identified from the predicted gene coding sequences using the Roary 3.6.0^86^ at 90% identity threshold. The pan-genome and core-genome accumulation curves were computed with 10 iterations. The gene presence-absence table was generated by combining Prokka^85^ and Roary^86^ identified gene coding sequences and identifiers. Most pan- and core-genome genes were shared across species; the largest difference between the number of core genes (∼2,000 per species) and accessory/pangenome genes (∼3,000–6,000 per species) within species indicates substantial strain-level differences in gene content between *Staphylococcus* (Figures S5A-S5C).

### Functional gene annotation and identification of differential features between species and clusters

COG functional categories were annotated using eggNOG-mapper 2.1.12^87^ with -i diamond. KEGG orthologs of each gene were annotated by blasting gene sequences to a customed KEGG ortholog database UBLAST (USEARCH v8.0.1517^88^) at e-value threshold of 10^9^. Identified KEGG ortholog numbers were further mapped to a customed KO identifier-KEGG mapping datasheet for annotation of KEGG ortholog, KEGG reaction, KEGG pathway, and COG category. The representative KEGG ortholog number and COG category of each gene was determined by majority rule. KEGG module completeness analysis was performed by MicrobeAnnotator v2.0.5 with -m diamond.

BGCs were identified using antiSMASH 7.0.0^89^ with default parameters. Virulence factors were annotated by blasting gene sequences against *Staphylococcus*-specific genes in VFDB^90^ with the addition of four phenol-soluble modules (PSMs)^91^ using UBLAST (USEARCH v8.0.1517^88^) at 50% identity threshold and expect value (e-value) threshold of 10^-9^, and five αPSMs^92^ at 60% identity threshold and e-value threshold of 10.

Differential gene content analysis for VFs, BGCs, KOs and genes between species and clusters was performed by MaAsLin2^80^ using a formula of gene content presence/absence matrix ∼ species + cluster. *S. hominis* and Commen_c2 were chosen as reference level for species and cluster.

### Identification of strain-level differential genes of *S. epidermidis*

*S. epidermidis* strain-level gene presence/absence was determined from genome annotations as described above. Strains that significantly upregulated innate immunity genes in RHE were assigned to the DEG group, while those that did not were assigned to the non-DEG group. For each innate immunity gene, the association between the presence of each *S. epidermidis* gene and strain group (DEG vs. non-DEG) was evaluated using 2×2 contingency tables. Statistical significance was assessed using Fisher’s exact test, with multiple testing correction performed using the Benjamini–Hochberg method.

### Transcriptional profiling of *Staphylococcus* transcriptome

The STAR^73^ filtered reads that were unmapped to human CHM13v2.0 reference genome were mapped to corresponding assembled genome of *Staphylococcus* to obtain corresponding bacterial reads using bowtie2 2.4.5^93^. Then read count aligned to each gene or ribosomal RNA gene was computed using featureCounts v1.6.4^74^ (from Subread1.6.4) with -t CDS or -t rRNA, respectively. The reads were averaged by triplicates, and TPM of each gene was calculated. On average, ∼2.7 million microbial reads per sample were retrieved. ∼100K m RNA reads per sample were retrieved, corresponding to ∼40X genome coverage and resulting in ∼75% of genes analyzable for >80% of strains (Figures S6A-S6E).

### Correlation analysis between *Staphylococcus* transcriptome and RHE gene module

Genes present in over 50% of *Staphylococcus* genomes and expressed in over 80% strains that have the gene (referred as prevalent genes) were chosen for correlation analyses, which were performed between the Z-scored normalized expression of prevalent genes of *Staphylococcus* and RHE gene module scores by HAllA 0.8.20^94^.

### High-resolution microbial metabolomic profiling

Each strain within collection of *Staphylococcus* strains were grown to saturation in triplicate in a single batch of rich media (TSB+vitamin K+hemin) to control for potential media batch effects, then normalized to 100 million cells by measuring culture optical density (OD600) and total cell counts using ultra-rapid flow cytometry (iQUE screener) and SYTO-based dye (Invitrogen). Undiluted supernatants and normalized microbial cell pellets are snap-frozen and stored at −80°C until assayed. Microbial supernatants and cell pellets were extracted using established protocols^95^. Extracts were aliquoted, dried, and reconstituted for ultra-high performance liquid chromatography (UHPLC)-mass spectrometry (MS) metabolomics, using a hydrophilic interaction (HILIC) column with positive electrospray ionization (ESI) and a reversed-phase C18 column with negative ESI, as per our previous work^96–98^. MS/MS data was collected on pooled samples, i.e., a mixture of all or representative biological samples. In-house chemical libraries (∼1100 chemicals) were used to annotate metabolites. The high-throughput samples were profiled singly since the analytical reproducibility on this platform was validated with rigorous QA/QC protocols. Data was processed using XCMS and optimal peak extraction parameters were determined empirically. MS/MS data was searched against *asari*^99^, mzCloud, MoNA, METLIN^100^, and HMDB^101^ for additional annotation. Metabolites on the arginine pathway were annotated. A more detailed annotation of the extensive unknowns in this dataset is ongoing in a separate endeavor.

### Analysis of similarity and distance between *Staphylococcus* strain pairs between multiple omics

*Staphylococcus* genomic, metabolic and transcriptomic similarities between *Staphylococcus* strain pairs and similarities of transcriptomic responses of RHE to *Staphylococcus* were calculated from *Staphylococcus* gene presence/absence, normalized metabolic feature abundance, TPM and RHE log2FC matrices, respectively. Correlation analyses were applied to the following combinations: *Staphylococcus* genomes & metabolomes, *Staphylococcus* genomes & transcriptomic responses of RHE to *Staphylococcus*, and *Staphylococcus* & transcriptomic responses of RHE to *Staphylococcus*. Distances were then inferred by (1-Pearson’s correlation coefficient) for each *Staphylococcus* strain pair.

### Biofilm formation assay of *Staphylococcus* strains

Biofilm formation was evaluated using a modified crystal violet staining protocol based on previously published methods^102,103^. A set of inflammatory *S. epidermidis* strains (n = 25), commensal *S. epidermidis* strains (n = 12), positive and negative control strains of known biofilm-formers (*S. aureus* USA300LAC, *S. epidermidis* 1457 (Sep_1457)) and low biofilm formers (*S. epidermidis* ATCC12228 (Sep_ATCC12228)) were examined. *Staphylococcus* strains were streaked from frozen glycerol stocks and inoculated into 3 mL of TSB (Becton, Dickinson and Company; #211825) and grown overnight at 37°C with shaking. Overnight cultures were measured for optical density and diluted 1:100 in TSB supplemented with 1% glucose to enhance biofilm formation. 150 µL of the diluted cultures were added in technical replicates (n=3-4) to 96-well plates, with wells containing 150 µL of sterile TSB + 1% glucose served as negative controls. Plates were incubated statically at 37°C for 16 hours, then culture supernatant was aspirated. Biofilm presence was confirmed by visually inspecting and by measuring optical density (OD) 600 of the supernatants was taken to confirm bacterial growth. Biofilms were washed twice with 200 µL of sterile deionized water. Plates were then heat-fixed at 60°C for 30 minutes. Biofilms were stained with 100 µL of 0.1% crystal violet (Sigma-Aldrich; #C3886) for 10 minutes at room temperature. Excess stain was aspirated, and wells were washed twice with 200 µL of sterile deionized water. Crystal violet retained by the biofilms was solubilized in 200 µL of 70% isopropanol and incubated for 30 minutes at room temperature. 150 µL of the solubilized stain was transferred to a new 96-well plate, and absorbance was measured at OD595 using a plate reader (BioTek Synergy H1, Agilent Technologies). Each strain was assayed independent experimental replicates (n=3) performed on separate days.

### Blood survival assay of *Staphylococcus* strains

Survival ability of *Staphylococcus* strains in rabbit blood was assessed. A set of inflammatory *S. epidermidis* strains (n = 7), commensal *S. epidermidis* strains (n = 6), and a positive control of a known bloodstream infection strain (*S. aureus* USA300LAC) were examined. For the complement depletion assay, a subset of inflammatory *S. epidermidis* strains (n = 3), commensal *S. epidermidis* strains (n = 3), and *S. aureus* USA300LAC) of above were examined. *Staphylococcus* strains were streaked from frozen glycerol stocks and inoculated into 3 mL of TSB (Becton, Dickinson and Company; #211825) and grown overnight at 37°C with shaking. Overnight cultures were pelleted, washed once with sterile PBS, and resuspended to an optical density at 600 nm (OD_600_) of 1.0, corresponding to approximately 1 × 10⁸ CFU/mL for each strain.

Rabbit red blood cells with complement (Innovative Research Inc.; #IRBRBC10ML) was aliquoted into 96-well plates (150 µL per well). Bacteria were added to blood to achieve a final inoculum of approximately 1 × 10^6^ CFU per well. For complement depletion assay, blood was centrifuged to separate plasma from cellular components. Plasma was collected and heat-inactivated at 56°C for 20 minutes, while the remaining red blood cell fraction was washed once with 1 mL of sterile PBS and centrifuged at 3,500 × g for 10 minutes to remove residual plasma. Heat-inactivated plasma was then recombined with the washed cellular fraction to reconstitute complement-depleted blood. Inoculum controls were plated to determine the exact number of bacteria added at 0h. Plates were incubated at 37°C with continuous shaking. Following incubation, samples were serially diluted in sterile PBS to determine CFU at 2h or 24h. Each strain was assayed with biological replicates (n=2) and technical replicates (n=2).

### Growth assay of *S. epidermidis* strains in defined media

Growth kinetics of *S. epidermidis* strains were assessed using a microplate-based assay under arginine-supplemented and arginine-free defined media adapted from (^45^ and ^104^). A set of inflammatory *S. epidermidis* strains (n = 18) and commensal *S. epidermidis* strains (n = 10) were selected for the experiment. Strains were streaked from frozen glycerol stocks and inoculated into 3 mL of TSB (Becton, Dickinson, and Company; #211825), followed by overnight incubation at 37°C with shaking. 1 mL of each overnight culture was pelleted by centrifugation, washed once with 1 mL of sterile PBS, pelleted again, and resuspended in 1 mL PBS.

For inoculation, washed bacteria were diluted 1:100 in defined media with or without arginine supplementation. The composition of the defined media is described in Table S7. Each condition was plated in quadruplicate (4 × 200 µL) in a single 96-well microplate, and each strain was tested in two independent experiments, resulting in eight replicates per strain. Growth was monitored over time by measuring optical density (OD600) at 0.5-hour intervals for 20 hours using a microplate reader (BioTek Synergy H1, Agilent Technologies). Growth metrics were calculated using gcplyr^105^, including maximum growth rate, maximum density and AUC of growth curve.

### Validation cohort analysis to demonstrate robustness of cluster classifications

To validate the robustness of our findings, we conducted two subsequent validation experiments with a subset of strains as described above. A KNN classification model was first trained on commensal/inflammatory commensal clusters defined by transcriptional responses of responsive innate immunity genes of RHE to *Staphylococcus* colonization in the original 180-strain experiment using scikit-learn^106^ Python package. Then the commensal/inflammatory commensal cluster labels of transcriptional responses of RHE to each of the 9 *Staphylococcus* strains in the validation experiment was predicted by the KNN model above. If the predicted cluster label was the same with the strain’s cluster label in the original dataset, then it was marked a correct classification. *S. pasteuri* was classified by the same KNN model above as well to predict its commensal cluster classification.

### Statistics and sample comparisons

Data was analyzed and visualized using the following Python packages: seaborn 1.11.3^81^, scikit-learn 1.3.2^106^, statannot 0.2.3, pandas 2.1.2^107^, numpy 1.26.1^108^, and matplotlib 3.7.3^109^; and R packages: ComplexHeatmap 2.25.1^110^, and ggplot2 3.5.1.

## DATA AND CODE AVAILABILITY

Raw RNA sequencing data related to this article can be accessed in National Center for Biotechnology Information dbGaP accession number phs004648. Original code for analysis framework of this study is available at https://github.com/ohlab/Staph-RHE.

## ACKNOWLEDGEMENTS AND FUNDING

We are thankful to the Oh laboratory for inspiring discussions and acknowledge the contribution of the Genome Technologies Service at The Jackson Laboratory and Sequencing and Genomic Technologies at Duke University School of Medicine for expert assistance with sample sequencing for the work described in this publication. Funding: National Institutes of Health 1 DP2 GM126893-01 (JO), 5 R21 AR075174 (JO), 7R01AR083742 (JO), 7R21AR082668-02 (JO), LEO FOUNDATION GRANT #LF-OC-24-001564 (JO).

## AUTHOR CONTRIBUTIONS

Conceptualization was carried out by JO and LMM. Data curation was performed by RY, MMS and ESA. Formal analysis was conducted by RY. Investigation was carried out by RY, MMS, ESA, WZ, AYV., GMW, AYK, MT and MG. Project administration was handled by JO and LMM. Resources were provided by JO, LMM and SL. Funding acquisition was secured by JO and LMM. Supervision was provided by JO and LMM. Visualization and writing – original draft were completed by RY. Writing – review and editing were performed JO and LMM.

## SUPPLEMENTARY FIGURES

**Figure S1.**
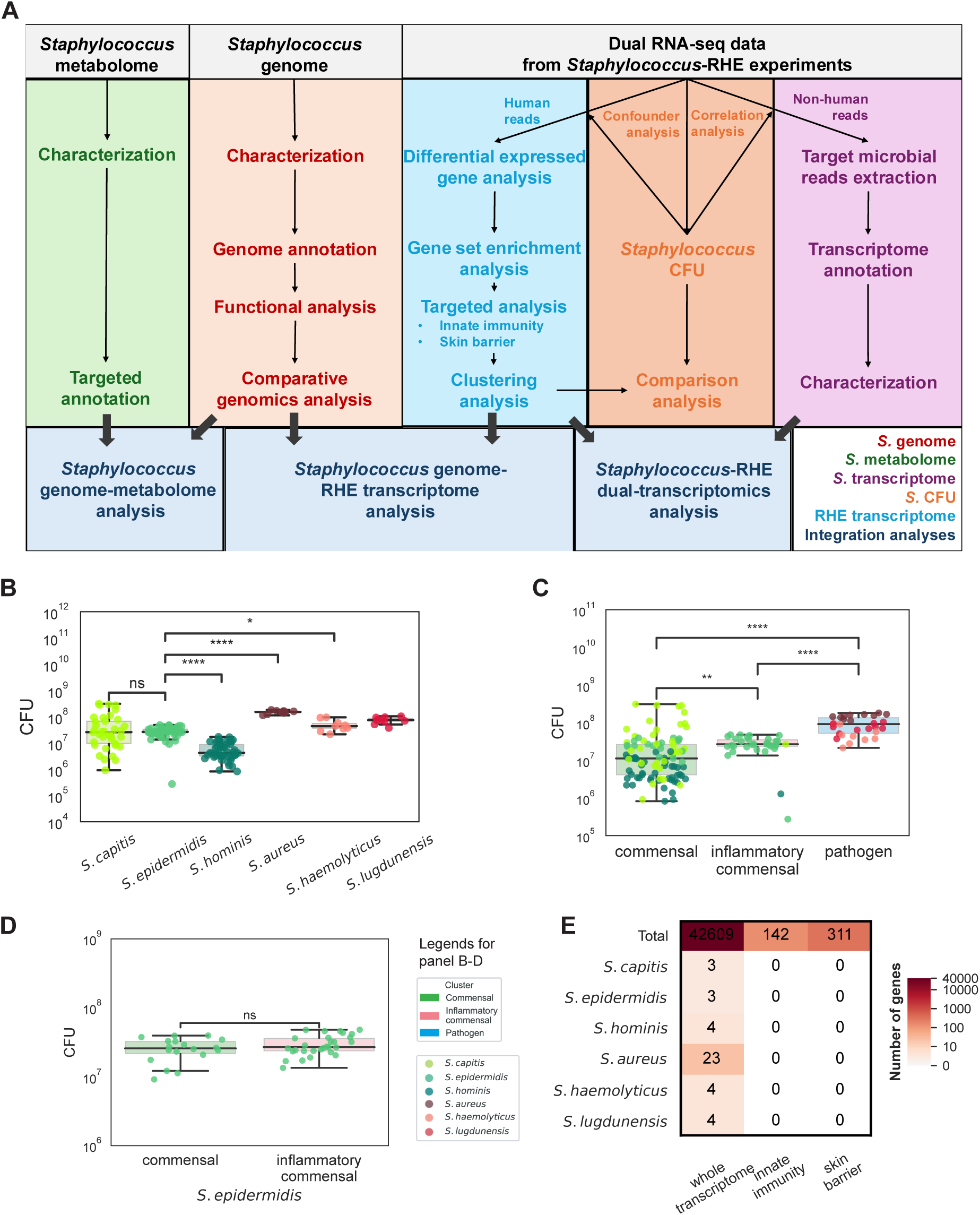
Analysis overview; confirmation that *Staphylococcus* growth does not explain RHE transcriptional responses, related to **Figure 1 and 2**. **(A)** Scheme of analyses. Multiple ‘omics datasets include dual RNA-seq data of *Staphylococcus* (*S.*) and reconstructed human epidermis (RHE) following an 18-hour colonization, *Staphylococcus* genomes, and untargeted metabolomics of *Staphylococcus* cell pellets and supernatants. Integrated analyses were performed across datasets. **(B-D)** *Staphylococcus* burden across conditions. Boxplots of colony-forming units (CFUs) grouped by species (B), cluster (C), or *S. epidermidis* only (D) at 18 hours. Points represent mean CFUs per strain (n = 3 replicates), colored by species or cluster. Boxes represent IQR with median. **** p ≤ 0.0001, ** 0.001 < p ≤ 0.01, * 0.01 < p ≤ 0.05., ^ns^ p > 0.05. **(E)** Genes in RHE associated with *Staphylococcus* burden. Genes whose expression correlated with CFUs were identified by MaAsLin2 linear regression (FDR-adjusted P ≤ 0.05). Rows indicate confounded genes across the whole transcriptome, innate immunity panel, or skin barrier panel.

**Figure S2.**
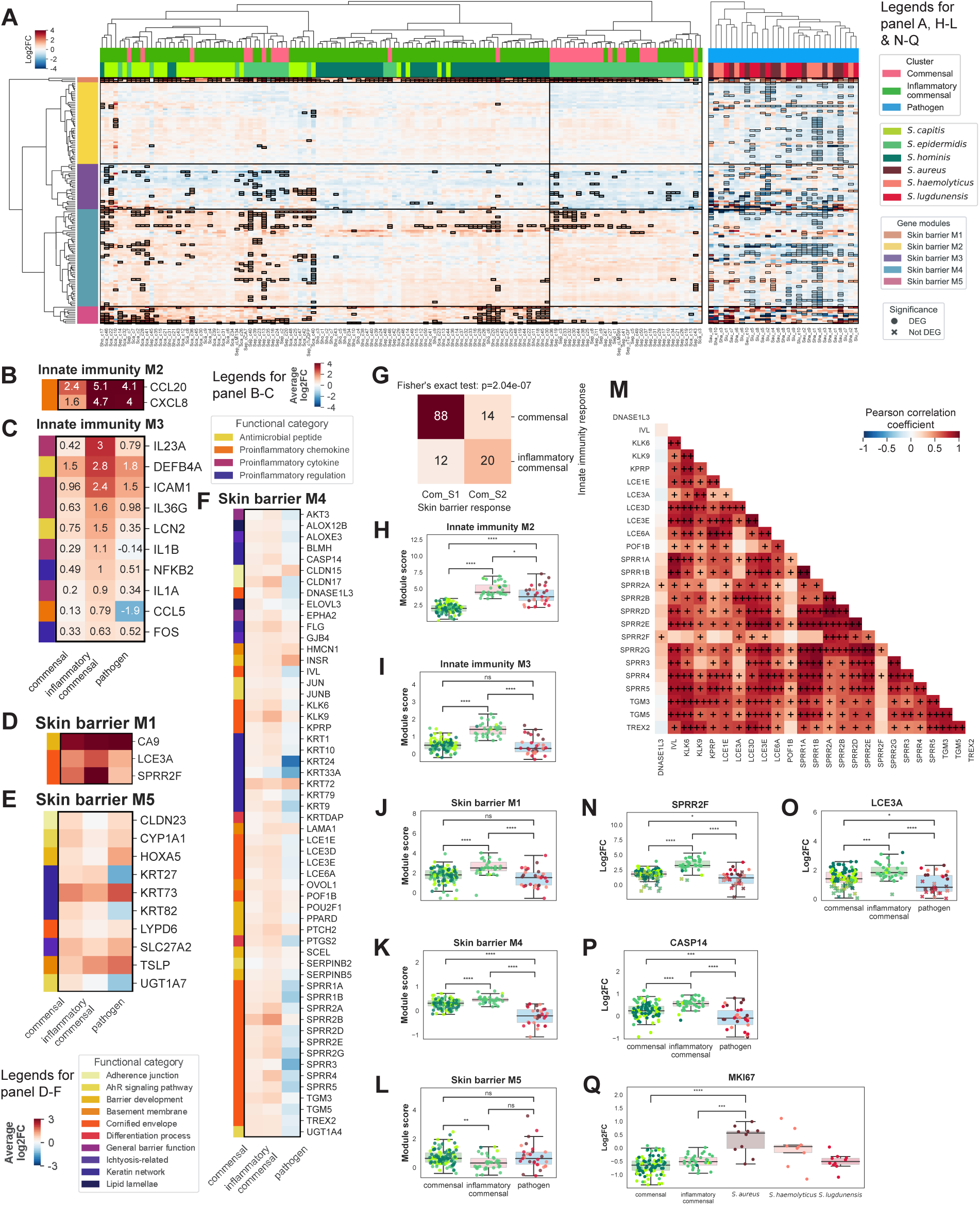
Differential innate immunity and skin barrier responses in RHE to *Staphylococcus* colonization, related to Figure 2. **(A)** Heatmap of skin barrier gene expression (log2 fold-change (FC)) in RHE following colonization by *Staphylococcus* strains (n=3 replicates per strain) relative to vehicle controls (n=14 replicates). Columns represent strains; rows represent genes from a curated skin barrier panel. Outlined cells indicate statistical significance (FDR adjusted P-value ≤ 0.05 and |log2FC| ≥ 1). Strains are grouped into commensal, inflammatory commensal, and pathogen and colored by species. Genes are organized into modules (skin barrier M1-5) to reduce data dimensionality based on hierarchical clustering. **(B-F)** Heatmaps (log2FC) of module-level responses for innate immunity M2 (B), M3 (C), skin barrier M1 (D), M5 (E) or M4 (F) averaged by *Staphylococcus* cluster. Functional categories are indicated to the left. **(G)** Concordance of clustering methods based on innate immunity versus skin barrier responses. Statistical significance was evaluated using Fisher’s exact test. **(H-L)** Module scores (median log2FC) for innate immunity M2 (H), M3 (I), skin barrier M1 (J), M4 (K) or M5 (L). Points are colored by species and grouped by cluster as in A. **(M)** Correlations between cornified envelope genes enriched in skin barrier modules. Pearson correlation coefficients, ++, r ≥ 0.7; +, 0.3 ≤ r < 0.7; −, −0.7 ≤ r ≤ −0.3. **(N-Q)** Boxplots (log2FC) of representative barrier genes *SPRR2F* (N), *LCE3A* (O), or proliferative genes, *CASP14* (P), *MKI67* (Q). Points indicate statistical significance (FDR adjusted P-value ≤ 0.05 and |log2FC| ≥ 1 and colors are as in A). **** p ≤ 0.0001, *** 0.0001 < p ≤ 0.001, ** 0.001 < p ≤ 0.01, * 0.01 < p ≤ 0.05., ^ns^ p > 0.05.

**Figure S3.**
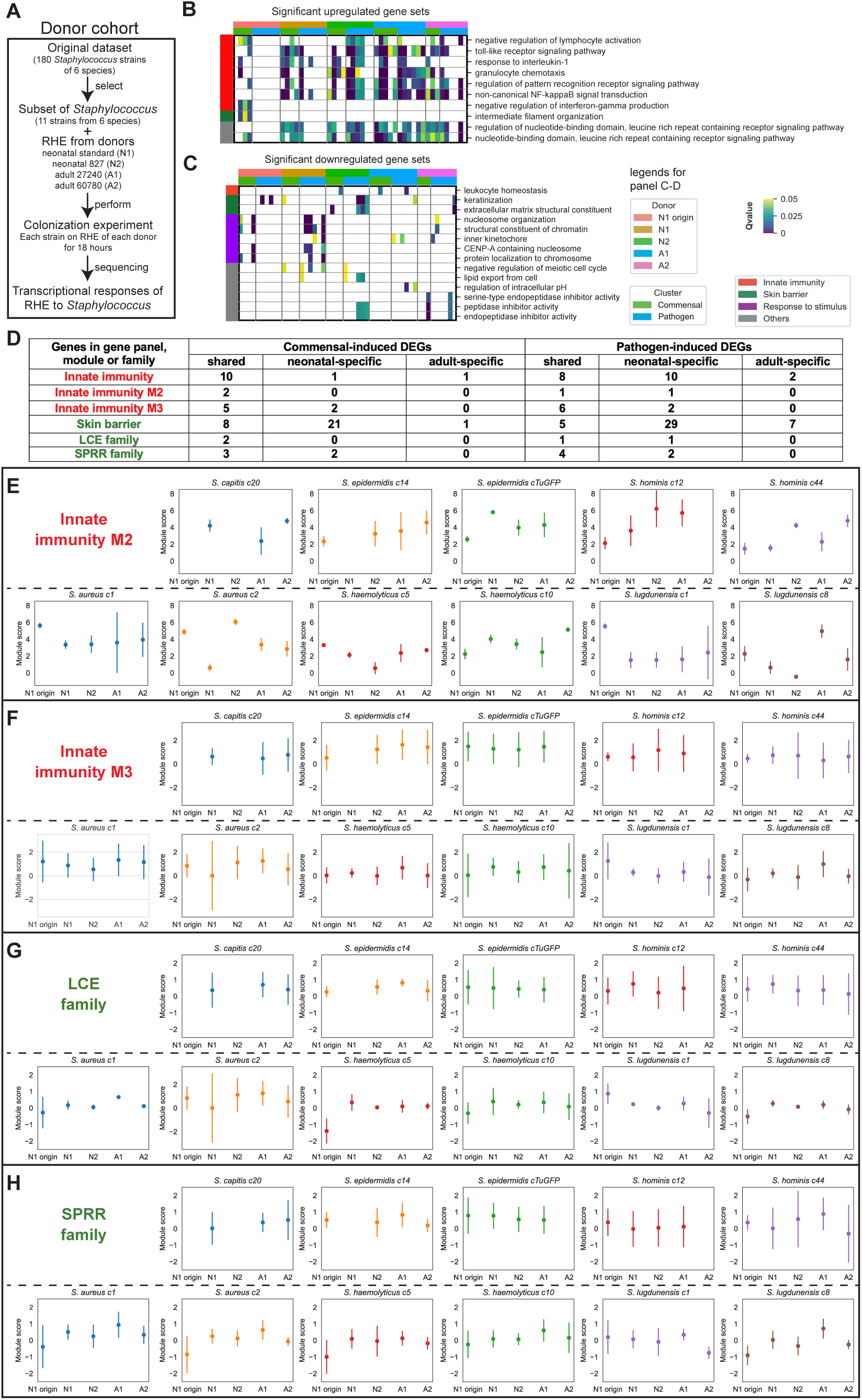
Transcriptional responses of RHE across keratinocyte donors are largely conserved, related to Figure 3. **(A)** Experimental design. 11 random strains from 6 *Staphylococcus* species were applied to RHE of 4 donors, two neonatal (standard “N1” and 827 “N2”) two adult (27240 “A1” and 60780 “A2”) and processed as previously. **(B-C)** Heatmaps of enriched Gene Ontology (GO) gene sets among upregulated (B) and downregulated genes (C) in *Staphylococcus-*colonized RHE relative to controls from both the original dataset and donor cohort. Rows represent GO gene sets, columns represent strains; cells indicate enrichment (Q ≤ 0.05) and nonsignificant entries are blank. |**(D)** Shared and donor-specific upregulated DEGs in RHE. Table summarizing overlap of across neonatal and adult donors within innate immunity or skin barrier gene panels, module or family. **(E-H)** Module scores (median log2FC) for innate immunity M2 (E), M3 (F), and skin barrier gene families LCE (G) and SPRR (H). Points represent module scores; error bars indicate standard deviation across genes within the module.

**Figure S4.**
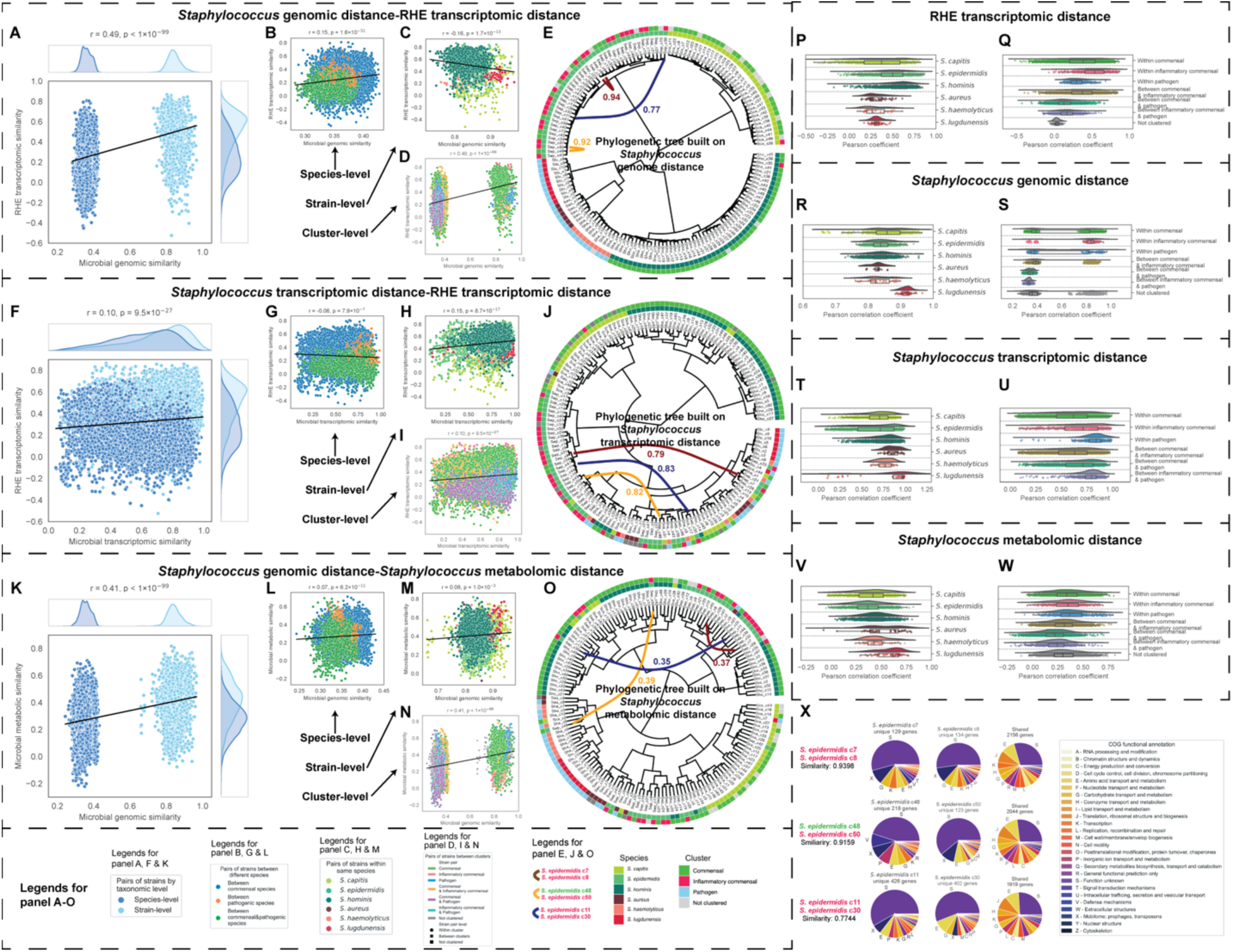
Genomic, transcriptomic and metabolomic differences in *Staphylococcus* associate with differential RHE responses, related to Figure 4, 5, 6 and 7. **(A-D)** Correlations between *Staphylococcus* genomic similarity and similarity of RHE transcriptional responses across strain pairs, shown globally (A) and stratified by species (B), strain (C), or cluster (D). Similarity was inferred by Pearson correlation coefficient based on gene presence–absence patterns or log2FC. Density plots indicate distribution of pairwise similarities. **(E)** Phylogenetic tree of *Staphylococcus* genomes and corresponding RHE transcriptome. **(F-I)** Correlations between *Staphylococcus* transcriptomic similarity and RHE transcriptional responses across strain pairs, globally (F), by species (G), strain (H), or cluster (I). Similarity was inferred by Pearson correlation coefficients of *Staphylococcus* strain pairs based on TPM-transformed *Staphylococcus* transcriptome. **(J)** Phylogenetic tree of *Staphylococcus* transcriptome and corresponding RHE transcriptome. **(K-N)** Correlations between *Staphylococcus* genomic and metabolomic similarity across strain pairs, globally (K), by species (L), strain (M), or cluster (N). Similarity was inferred by Pearson correlation coefficients of metabolic feature intensity within untargeted metabolomes. **(O)** Phylogenetic tree of *Staphylococcus* metabolome and corresponding genome. **(P-W)** Raincloud plots show distribution of pairwise intra-species and inter-cluster similarities of RHE transcriptomic responses (P-Q), *Staphylococcus* genomes (R-S), transcriptomes (T-U), and metabolomes (V-W). Distributions include a half-violin plot showing kernel density and boxplots (IQR with median). **(X)** Shared and unique genes in representative *S. epidermidis* strain pairs of note, e.g. highest similarity (c7&c8), highest inter-cluster similarity (c48&c50), or lowest similarity (c11&c30). Pie charts show genes by COG category.

**Figure S5.**
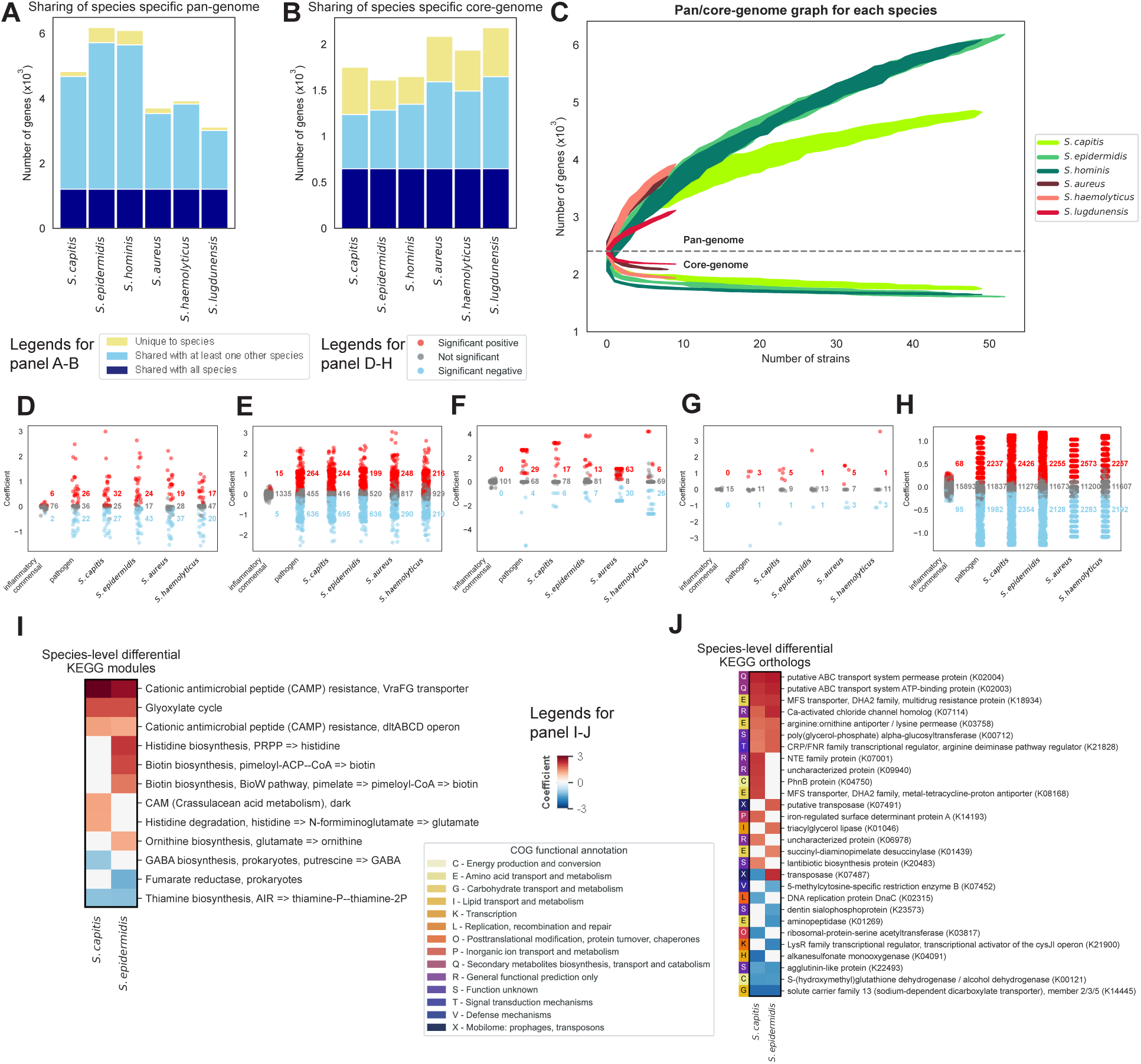
Species- and strain-level functional differences in *Staphylococcus* genomes, related to Figure 4 and 5. **(A-B)** Shared and unique species-specific pan (A) or core (B) genome across *Staphylococcus* species. **(C)** Gene accumulation curves for pan- and core genomes by species. Curves are colored by species; error bars show the standard deviation (n= 10 permutations). **(D-H)** Differential genomic features between *Staphylococcus* clusters or species, including KEGG modules (D), KEGG orthologs (E), virulence factors (VFs) (F), biosynthetic gene clusters (BGCs) (G), and genes (H). Significant features were identified by linear regression (FDR adjusted P-value ≤ 0.05). Columns indicate clusters or species; values indicate feature counts (sum of gene contents/category). **(I-J)** Representative differential KEGG modules (I) or orthologs (J) across species. COG functional category is indicated at the left.

**Figure S6.**
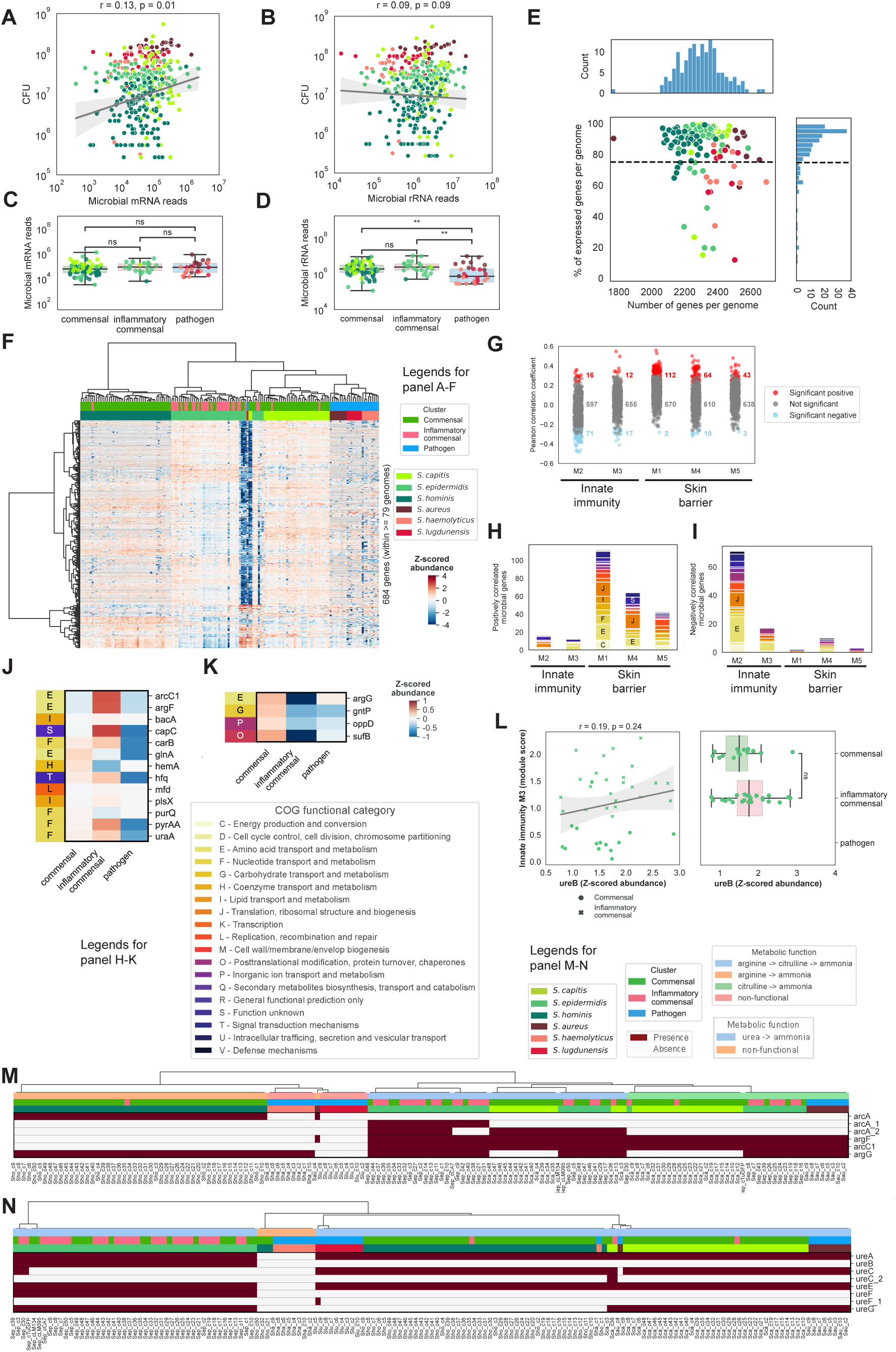
*Staphylococcus* transcriptional differences associate with differential effects on RHE, related to Figure 6. **(A-B)** Correlation between *Staphylococcus* mRNA (A) or rRNA (B) abundance and CFUs during RHE colonization. Points represent mean values (n=3); Pearson correlation coefficients (r), fitted regression line, and P-values. **(C-D)** Comparisons of *Staphylococcus* mRNA (C) or rRNA (D) reads across clusters. ** 0.001 < p ≤ 0.01, ^ns^ p > 0.05. **(E)** Fraction of expressed genes detected per *Staphylococcus* genome (y-axis) with respect to total number of genes (x-axis). Histograms at top and side show frequencies; dashed line shows cutoff of 0.75 that encompasses. **(F)** Expression of prevalent *Staphylococcus* genes (n=684), defined as present in ≥ 50% (79) genomes and detected in ≥ 0.75 of genes/genome. **(G)** Correlation between expression of prevalent *Staphylococcus* genes and RHE module scores (FDR adjusted P-value < 0.05). **(H-I)** COG functional categories of genes significantly positively (H) or negatively (I) correlated with RHE modules. **(J-K)** Expression of representative *Staphylococcus* genes positively (J) or negatively (K) correlated with RHE modules across clusters; COG categories are indicated at left. **(L)** Correlation between *Staphylococcus ureB* expression (TPM, x-axis) and RHE innate immunity module M3 (median log2FC, y-axis). Pearson correlation coefficient (r) and P-value (p) are indicated. Box plots indicate expression by cluster (IQR with median). ^ns^ p > 0.05. **(M-N)** Presence of genes in the ammonia-producing arginine pathway (M) or urea pathway (N). Cells of heatmap were colored in red if gene present in *Staphylococcus* genome.

**Figure S7.**
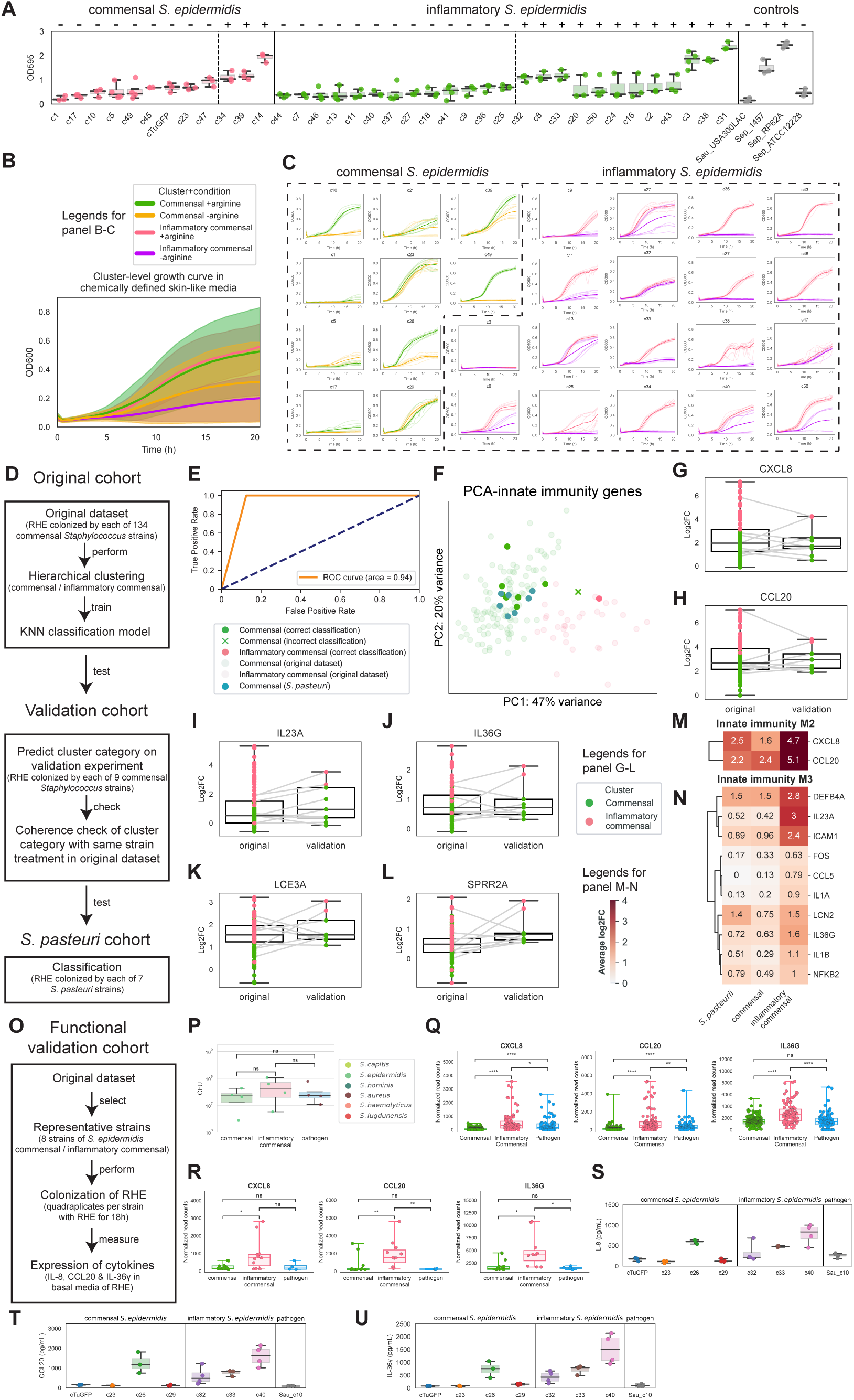
Strain-level metabolomic and phenotypic differences in *Staphylococcus* underlie their differential effects on RHE, related to Figure 7. (A) Biofilm-forming capacity of inflammatory vs. commensal *S. epidermidis* strains. (-) or (+) indicates negative or positive for biofilm by comparing to a canonical biofilm-forming strain (Sep_1457). Control strains have characterized biofilm formation (*S. aureus* USA300LAC (-), *S. epidermidis* 1457 (+), RP62A(+), and ATCC12228(-)). (B) **(B-C)** Growth curves of *S. epidermidis* strains in chemically defined media, averaged by cluster **(B)**or individual strain (C). For (B), error bars represent the standard deviation. **(D)** Workflow for validation of cluster assignments using other cohorts. A k-nearest neighbors (KNN) classifier was trained on the original dataset and applied to a new validation cohort of RHE colonized with 9 different strains. Classification of cluster was validated with *S. pasteuri* strains (*S. pasteuri* cohort, n = 7 strains) using the trained KNN model built. **(E)** ROC curve of classifier performance; area under curve (AUC = 0.94) indicates prediction accuracy. **(F)** PCA of RHE innate immunity gene responses in original dataset, validation and *S. pasteuri* cohorts. Points are colored by cluster or species and transparency the dataset origin. **(G-L)** Boxplots of representative gene expression (log2FC) in RHE for innate immunity genes *CXCL8* (G), *CCL20* (H), *IL23A* (I) and *IL36G* (J), and skin barrier genes *LCE3A* (K) or *SPRR2A* (**L**) comparing original and validation datasets. Lines connect matched strains. Boxes indicate IQR with median. **(M-N)** Heatmaps of innate immunity M2 (M) and M3 (N) gene expression with average log2FC) for *Staphylococcus* clusters or *S. pasteuri* strains. **(O)** Workflow for functional validation of *S. epidermidis* strains. RHE were colonized with representative inflammatory commensal (n=3), commensal (n=4), and pathogenic (n=1) strains. **(P)** CFUs at endpoint for this functional validation cohort do not explain cluster assignment, as in the original experiment. Points are average CFUs across n=3-4 replicates. ^ns^ p > 0.05. **(Q-R)** RNA expression of cytokines *CXCL8* (left), *CCL20* (middle) and *IL36G* (right) recapitulate results from the original experiment (Q) or functional validation cohort (R). **(S-U)** Protein expression as measured by ELISA of basal media following colonization for IL-8 (S), CCL20 (T) and IL-36γ (U)

## SUPPLEMENTARY TABLES

Supplementary Table S1. Metadata and sequencing statistics for RHE colonization experiments, related to Figure 1. Summary of sample information, strain identity, and sequencing depth metrics for dual RNA-seq datasets. Sheets include:

- *Staph_origin*: strain, species, cluster, experimental cohort and origin of *Staphylococcus* strains.
- *Dual-RNA-seq_stats*: sequencing depth, microbial reads and CFU.

Supplementary Table S2. RHE transcriptomic programs in response to *Staphylococcus* in 180-strain cohort, related to **Figures 1, 2 and 3**. Differential gene expression and gene set enrichment analyses of RHE transcriptomes in response to *Staphylococcus*. Sheets include:

- *Log2FC*: differential expression statistics for log2FC for genes in RHE.
- *Log2FC_significance*: significance of genes.
- *Shared_and_different_DEGs*: shared and unique DEGs.
- *GO_up_GeneRatio*: GO gene ratios for upregulated gene sets.
- *GO_up_qvalue*: adjusted p-values (q-values) for upregulated gene sets.
- *GO_down_GeneRatio*: GO gene ratios for downregulated gene sets.
- *Gene_down_qvalue*: q-values for downregulated gene sets.
- *Gene_lists*: gene panels for RHE in this study.

Supplementary Table S3. RHE transcriptomic programs in additional cohorts, related to Figure 3 and 7. Independent RNA-seq datasets used to validate key transcriptional changes observed in the primary analysis. Sheets include:

- *Donor_exp_log2FC*: differential expression statistics for log2FC for genes in RHE in donor cohort
- *Donor_exp_significance*: significance of genes in donor cohort.
- *Validation_exp_log2FC*: differential expression statistics for log2FC for genes in RHE in validation cohort.
- *Validation_exp_significance*: significance of genes in validation cohort.
- *Validation_exp_clustering*: clustering analysis of validation transcriptomes
- *Spasteuri_log2FC*: differential expression statistics for log2FC for genes in RHE for *S. pasteuri* cohort.
- *Spasteuri_significance*: significance of genes in *S. pasteuri* cohort.
- *Spasteuri_clustering*: clustering analysis for *S. pasteuri* cohort.

Supplementary Table S4. *Staphylococcus* genomic data, related to Figures 4 and 5. Gene contents information for S*taphylococcus* strains analyzed. Sheets include:

- *Gene_presence_absence*: gene presence/absence matrix.
- *Gene_annotation*: functional annotation of genes.
- *KEGG_module_completeness*: completeness of KEGG modules.
- *KEGG_ortholog*: KEGG orthologs.
- *Virulence_factors*: virulence factors.
- *Biosynthetic_gene_clusters*: predicted biosynthetic gene clusters.

Supplementary Table S5. *Staphylococcus* transcriptomic profiles, related to Figure 6. Expression profiles of *Staphylococcus* species derived from dual RNA-seq. Sheets include:

- *Staph_RNA-seq_TPM_averaged*: TPM values of genes of *Staphylococcus* averaged by strain.
- *Staph_top_genes_Zscored*: Z-score normalized expression of prevailing genes of *Staphylococcus*.

Supplementary Table S6. Multi-omics similarity analyses across *Staphylococcus* strains and RHE transcriptomic responses, related to Supplementary Figure S4.

Comparison of genomic, metabolic, and transcriptomic similarity across *Staphylococcus* strains and RHE transcriptomic responses. Sheets include:

- *Staph_genomic_similarity*: pairwise genomic similarities of each *Staphylococcus* strain pair.
- *Staph_metabolomic_similarity*: pairwise metabolomic similarities of each *Staphylococcus* strain pair.
- *Staph_transcriptomic_similarity*: pairwise transcriptomic similarities of each *Staphylococcus* strain pair.
- *RHE_transcriptomic_similarity*: pairwise transcriptomic similarities of RHE treated by each *Staphylococcus* strain pair.

Supplementary Table S7. *Staphylococcus* metabolomic and phenotypic profiles, related to Figure 7. *Staphylococcus* metabolomic profiles, all types of data for phenotypic assays, and results of validation cohorts. Sheets include:

- *Biofilm_formation*: quantification of biofilm formation capacity.
- *Blood_survival*: results of blood survival assays.
- *Blood_survival_comp_deplete*: results of blood survival assays under complement-depleted conditions.
- *Metabolomics_annotated*: annotated *Staphylococcus* metabolic profiles in this study.
- *CDM_ingredient*: ingredient for chemically defined media used in this study.
- *Growth_assay_SLM+arg*: results of growth assays under defined in chemically defined media with arginine.
- *Growth_assay_SLM-arg*: results of growth assays under defined in chemically defined media without arginine.
- *RHE_basal_media_ELISA*: results of ELISA of protein expression in basal media of RHE in functional validation cohort.

